# Microcephaly with a disproportionate hippocampal reduction, stem cell loss and neuronal lipid droplet symptoms in *Trappc9* KO mice

**DOI:** 10.1101/2023.11.20.567859

**Authors:** Sultan Aljuraysi, Mark Platt, Michela Pulix, Harish Poptani, Antonius Plagge

## Abstract

Mutations of the human *TRAFFICKING PROTEIN PARTICLE COMPLEX SUBUNIT 9* (*TRAPPC9*) cause a neurodevelopmental disorder characterised by microcephaly and intellectual disability. Trappc9 constitutes a subunit specific to the intracellular membrane-associated TrappII complex. The TrappII complex interacts with Rab11 and Rab18, the latter being specifically associated with lipid droplets (LDs). Here we used non-invasive imaging to characterise *Trappc9* knock-out (KO) mice as a model of the human hereditary disorder. KOs developed postnatal microcephaly with many grey and white matter regions being affected. *In vivo* MRI identified a disproportionately stronger volume reduction in the hippocampus, which was associated with a significant loss of Sox2-positive neural stem and progenitor cells. Diffusion Tensor imaging indicated a reduced organisation or integrity of white matter areas. *Trappc9* KOs displayed behavioural abnormalities in several tests related to exploration, learning and memory. Trappc9-deficient primary hippocampal neurons accumulated a larger LD volume per cell following Oleic Acid stimulation, and the coating of LDs by Perilipin-2 was much reduced. Additionally, *Trappc9* KOs developed obesity, which was significantly more severe in females than in males. Our findings indicate that, beyond previously reported Rab11-related vesicle transport defects, dysfunctions in LD homeostasis might contribute to the neurobiological symptoms of Trappc9 deficiency.

## Introduction

Intellectual disability and microcephaly are frequent symptoms of autosomal recessive neurodevelopmental disorders (Khan et al., 2016). Genetic screening of patients with such rare disorders has implicated an increasing number of genes with a variety of cellular functions (Khan et al., 2016). Mutations in *TRAFFICKING PROTEIN PARTICLE COMPLEX SUBUNIT 9* (*TRAPPC9*) were identified in patients with microcephaly, intellectual disability, inability to learn to speak and developmental delay (Koifman et al., 2010; Mir et al., 2009; Mochida et al., 2009; Philippe et al., 2009). Microcephaly in these patients was detectable by Magnetic Resonance Imaging (MRI) within the first year of life and included grey matter atrophies as well as white matter reductions (e.g. corpus callosum) (Amin et al., 2022; Aslanger et al., 2022; Ben Ayed et al., 2021; Bolat et al., 2022; Hnoonual et al., 2019; Koifman et al., 2010; Penon-Portmann et al., 2023; Radenkovic et al., 2022). Other disease symptoms occur more variably with obesity, dysmorphic facial features and hand-flapping movements being described in approximately half the patients (Aslanger et al., 2022; Bolat et al., 2022; Kramer et al., 2020).

Trappc9 forms a subunit of the metazoan TrappII multi-protein complex, which regulates vesicle trafficking, endosome recycling and lipid droplet homeostasis (Galindo & Munro, 2023; Li et al., 2017). TrappII and the related TrappIII complex share seven core subunits, but are distinguished by the association of specific subunits, i.e. Trappc9 and Trappc10 in TrappII and Trappc8, c11, c12 and c13 in TrappIII (Galindo & Munro, 2023). Recent structural analyses of the TrappII complex revealed a triangular shape with the large c9 and c10 subunits forming two sides of the triangle (Galindo & Munro, 2023; Jenkins et al., 2020). The complexes attach to intracellular membranes via binding to specific membrane-anchored Rab proteins, for which they have an activating guanine nucleotide exchange factor (GEF) function. The TrappII complex preferentially interacts with Rab11, Rab18, Rab19 and Rab43, but has also GEF activity for Rab1, which is however regarded as the main substrate for TrappIII (Galindo & Munro, 2023; Jenkins et al., 2020; Ke et al., 2020; Kiss et al., 2023; Li et al., 2017). Rab11 regulates endosome recycling and Trappc9-deficient neurons show a delay in recycling of the transferrin receptor (Ke et al., 2020). Rab18 is specifically associated with lipid droplets (LDs) and Rab18-as well as Trappc9-deficient cell lines and patients’ fibroblasts display enlarged LDs (Bekbulat et al., 2020; Carpanini et al., 2014; Deng et al., 2021; Li et al., 2017; Usman et al., 2022; Xu et al., 2018). Mutations in both, *RAB11B* and *RAB18*, cause neurodevelopmental disorders and intellectual disability with symptoms overlapping those of *TRAPPC9* deficiency (Bem et al., 2011; Carpanini et al., 2014; Cheng et al., 2015; Lamers et al., 2017). Furthermore, mutations in the various subunits of the Trapp complexes result in a number of distinct genetic disorders with partially overlapping phenotypes, termed ‘TRAPPopathies’, which hints at subunit-specific functions in addition to the functions of the full complexes (Sacher et al., 2019). For example, mutations in *TRAPPC10*, the other TrappII-specific subunit, causes symptoms similar to *TRAPPC9* deficiency, including microcephaly, corpus callosum thinning and intellectual disability, but obesity is absent in *TRAPPC10* patients (Rawlins et al., 2022).

The roles of Trappc9 and Rab18 in the cellular regulation of LDs has been little investigated so far for its potential relevance to the neurobiological symptoms of the disorders. On the other hand, recent work has revealed the importance of LDs in neural cells. LDs are generated at specialised sites of the endoplasmic reticulum, contain neutral lipids and cholesterol esters, are surrounded by a phospholipid monolayer and associated with various coat proteins (Olzmann & Carvalho, 2019; Sztalryd & Brasaemle, 2017). Neural stem and progenitor cells (NSPCs) of the subventricular zone and the dentate gyrus contain substantial numbers of LDs, and lipid metabolism as well as the amount of lipids stored influence their proliferative capability as well as differentiation tendencies (Ramosaj et al., 2021). Furthermore, Oleic Acid (OA) was recently identified as an important factor that stimulates NSPC proliferation through binding as an endogenous ligand to the nuclear receptor TLX/NR2E1 (Kandel et al., 2022). The activated receptor stimulates the expression of a range of cell cycle and neurogenesis genes in NSPCs (Kandel et al., 2022). Neurons generally contain fewer LDs, have a low capacity for fatty acid metabolism and are prone to lipotoxicity due to activity-induced fatty acid peroxidation (Ioannou, Jackson, et al., 2019; Ramosaj et al., 2021). However, the importance of regulation of lipid metabolism and LDs in neurons is demonstrated by two forms of hereditary spastic paraplegia. Mutations in the triglyceride hydrolase *DDHD2* are the cause for subtype SPG54 and mice deficient for Ddhd2 show LD accumulations in the brain, specifically in neurons but not glial cells (Inloes et al., 2014; Inloes et al., 2018). Furthermore, mutations in the Troyer syndrome gene *SPARTIN* lead to impaired autophagy of LDs and increased LD numbers in neurons (Chung et al., 2023). Despite these observations, our current mechanistic understanding of the pathogenic role of LD accumulations in neurons remains limited.

To investigate the neurodevelopmental disease mechanisms caused by *Trappc9* deficiency, we utilised a KO mouse line. *Trappc9* is located within the *Peg13-Kcnk9* cluster of imprinted genes on mouse chromosome 15 and human chromosome 8 (Ruf et al., 2007; Smith et al., 2003). Genomic imprinting is defined as parent-of-origin specific gene expression caused by epigenetic modifications that are inherited through either the maternal or paternal germline and maintained in somatic cells of the offspring (Ferguson-Smith, 2011; Tucci et al., 2019). Although genomic imprinting of *Peg13* and *Kcnk9* is conserved between mouse and human, this is not the case for the three neighbouring genes *Trappc9*, *Chrac1* and *Ago2* (Court et al., 2014), which are imprinted in murine brain tissue only where they show a ∼70 % expression bias from the maternal allele (Claxton et al., 2022; Liang et al., 2020; Perez et al., 2015). To focus on the strongest phenotypes, we have limited our analysis in this study to homozygous mutant mice, while a recent report reported that heterozygotes with a maternally inherited mutation (m-/p+) display symptoms almost as severe as homozygote KOs, while paternal heterozygotes (m+/p-) resemble wildtype mice (Liang et al., 2020). We show by *in vivo* MRI that the microcephaly develops postnatally and affects grey as well as white matter regions. We find that all analysed brain sub-regions are significantly smaller with the hippocampus being disproportionately more severely reduced relative to total brain volume. *Trappc9* is prominently expressed in hippocampal neurons as well as in adult NSPCs of the dentate gyrus with KO mice displaying reduced numbers of Sox2-positive NSPCs in this hippocampal sub-region. White matter in the corpus callosum (CC) shows less integrity as indicated by Diffusion Tensor Imaging (DTI). The microcephaly is accompanied by behavioural deficits related to cognition and learning in several test paradigms. On a cellular level, we provide further evidence for a role of Trappc9 in LD regulation as primary hippocampal KO neurons accumulate larger LD volumes in culture with reduced LD surface coating by Perilipin-2 (Plin2). Increased LDs are also present in adipose tissues of KOs, which display sex-specific differences in obesity.

## Results

### *Trappc9*-deficient mice develop microcephaly postnatally and show a disproportionate reduction in hippocampus volume

To investigate the mechanisms of disease caused by *TRAPPC9* mutations, we utilised the knock-out first model (Skarnes et al., 2011) of the International Mouse Phenotyping Consortium (IMPC), which has a gene trap cassette located in intron 5. We confirmed a lack of Trappc9 protein by Western blot of brain tissue from homozygous KO mice (Figure 1–figure supplement 1A). However, the tm1a LacZ-gene trap cassette of the targeted mutation did not result in any detectable β-Galactosidase expression from the *Trappc9* locus, which might be due to a cryptic splice donor site within the Engrailed2 component of the gene trap cassette, which we identified in RT-PCR products from KO brain and which resulted in splicing in and out of the En2 part and further onto *Trappc9* exon 6, thus disrupting the open reading frame (Figure 1–figure supplement 1B).

**Figure 1.**
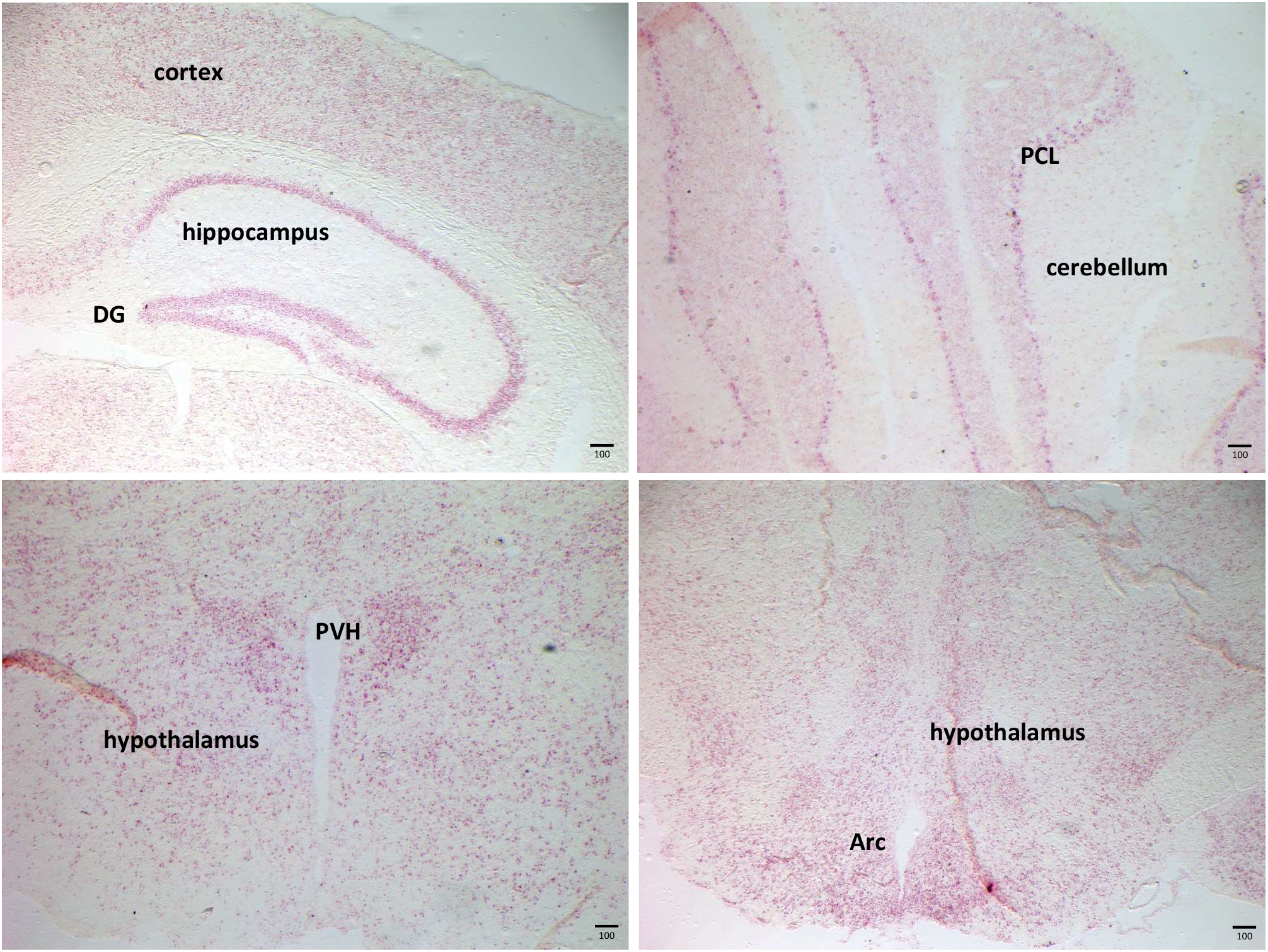
*Trappc9* expression pattern in the adult mouse brain. *In situ* hybridisation on coronal sections using RNAscope® probes indicates expression in many brain areas, including the cortex, hippocampus with dentate gyrus (DG), cerebellum with Purkinje cell layer (PCL) and hypothalamus with paraventricular nucleus (PVH) and arcuate nucleus (Arc). Scale bar: 100 µm.

To determine the *Trappc9* expression pattern in the adult mouse brain we used RNAscope *in situ* hybridisation probes and found widespread expression in many brain areas (Figure 1). The highest levels were detected in neuronal cell layers, i.e. the CA and granule cell layers of the hippocampus and dentate gyrus, Purkinje cell layer of the cerebellum as well as the hypothalamic paraventricular and arcuate nuclei. These findings are in line with the Allen Brain Map single-cell RNA expression data, which indicate highest levels of *Trappc9* in various types of cortical and hippocampal neurons while much lower levels have been detected in astrocytes and oligodendrocytes (https://portal.brain-map.org/atlases-and-data/rnaseq).

One of the most consistent symptoms of patients with *TRAPPC9* mutations is microcephaly, which has been detected in children as early as one year of age and includes reduced white matter (e.g. corpus callosum), cerebral and cerebellar atrophies (Amin et al., 2022; Aslanger et al., 2022; Ben Ayed et al., 2021; Bolat et al., 2022; Hnoonual et al., 2019; Koifman et al., 2010; Penon-Portmann et al., 2023; Radenkovic et al., 2022). To investigate microcephaly in homozygous *Trappc9* mutant mice, we determined brain volumes via MRI as well as tissue weights at birth, weaning and adult stages. We found no difference in brain volumes or weights on the day of birth (Figure 2A, B, Tables 1 and 2). However, we observed significantly smaller brain volumes in KO mice at weaning age, which was in line with tissue weight differences (Figure 2A, B; Tables 1, 2). These data indicate a postnatally developing microcephaly of 6 – 7 % by weaning age. To assess the microcephaly phenotype longitudinally across age, we acquired *in vivo* MRI scans in a cohort of mice at young (14 – 18 weeks) and mature adult stages (40 – 42 weeks). Genotype, but not age, had a significant effect on brain volume at both time points (Figure 2A; Table 1). Brain weight data confirmed an 8 – 10 % reduction in adult KO mice of both sexes (Figure 2B; Table 2).

**Figure 2:**
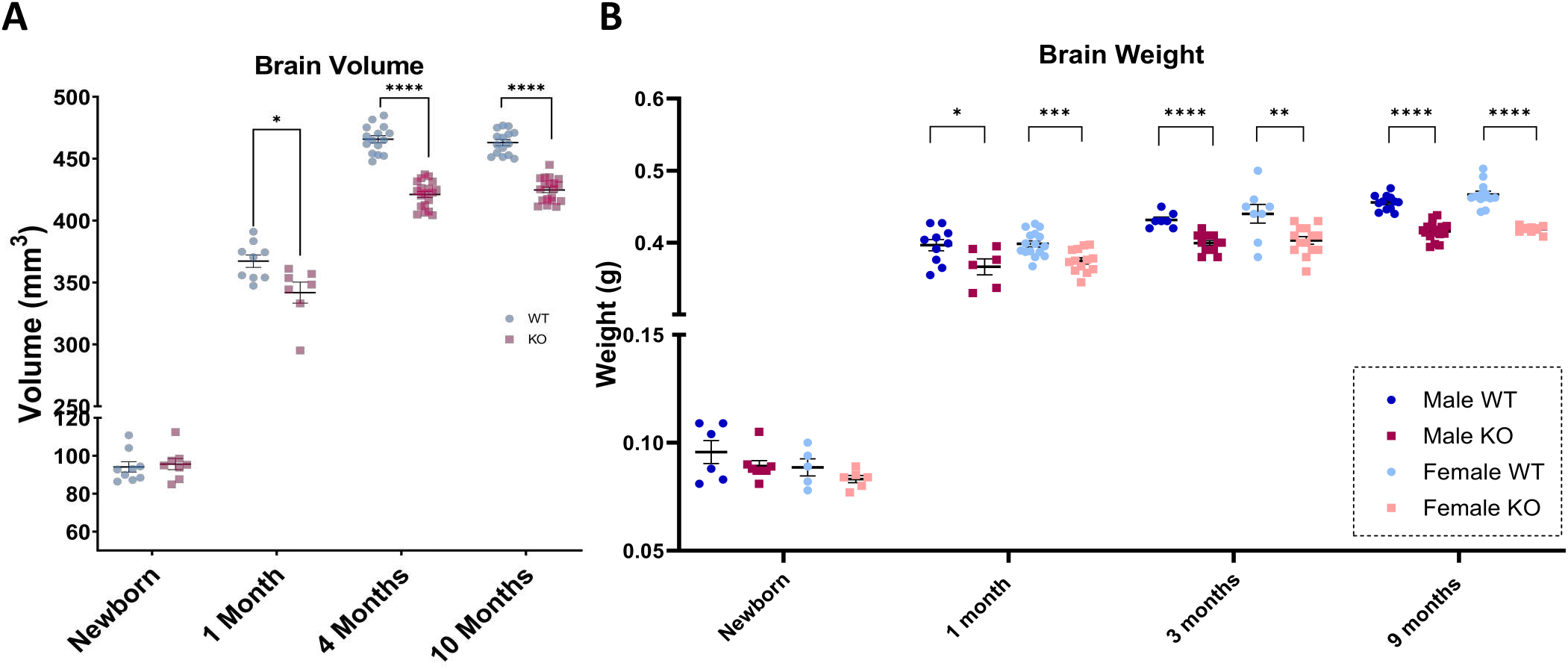
*Trappc9* deficient mice develop postnatal microcephaly. **A)** Whole brain volumes measured by MRI. Data for males and females of the same genotype were pooled, since no difference was found between sexes. Data for newborn and 1-month old mice are based on *ex vivo* T1-weighted imaging of whole skulls to avoid brain dissection artefacts (independent *t.*test, p: *<0.05). Data for 4- and 10-months old mice are based on *in vivo* longitudinal T2-weighted imaging of the same cohort of mice (two-way repeated measures ANOVA with Šídák’s multiple comparison test, p: ****<0.0001). Table 1. **B)** Brain tissue weights of *Trappc9* WT and KO males and females at various ages (independent *t.*test, p: *<0.05, **<0.01, ***<0.001, ****<0.0001). Data presented as mean ± sem. Table 2.

**Table 1.** Brain volumes of WT and *Trappc9* KO mice at different ages as measured by MRI. The sample size (N) is given in brackets for each group; statistical comparisons by independent *t.*test (newborn, 1 month) or two-way repeated measures ANOVA with Šídák’s multiple comparison test (4 and 10 months).

| Brain Volume (mm<sup>3</sup>), mean ± sem |  |  |  |  |
| --- | --- | --- | --- | --- |
| Age | WT | KO | Relative difference to WT | p-value |
| Newborn | 94.1 ± 2.74 (9) | 95.6 ± 2.93 (8) | +1.59 % | 0.71 |
| 1 month | 367.3 ± 5.02 (9) | 341.9 ± 8.51 (7) | -6.92 % | 0.017 |
| 4 months | 465.8 ± 2.84 (15) | 421.1 ± 2.47 (19) | -9.60 % | <0.0001 |
| 10 months | 463.1 ± 2.49 (15) | 424.8 ± 2.24 (19) | -8.27 % | <0.0001 |

**Table 2.** Brain weights of WT and *Trappc9* KO mice at different ages. The sample size (N) is given in brackets for each group; statistical comparisons by independent *t*.test.

| Brain weight (g), mean ± sem |  |  |  |  |  |  |  |  |
| --- | --- | --- | --- | --- | --- | --- | --- | --- |
|  | Male |  |  |  | Female |  |  |  |
| Age | WT | KO | Relative difference to WT | p-value | WT | KO | Relative difference to WT | p-value |
| Newborn | 0.096 ± 0.005 (6) | 0.089 ± 0.002 (8) | -7.3 % | 0.26 | 0.089 ± 0.004 (5) | 0.083 ± 0.002 (6) | -6.7 % | 0.21 |
| 1 month | 0.396 ± 0.008 (10) | 0.366 ± 0.011 (6) | -7.6 % | 0.039 | 0.398 ± 0.004 (16) | 0.375 ± 0.004 (13) | -5.8 % | 0.0007 |
| 3 months | 0.431 ± 0.004 (7) | 0.399 ± 0.003 (13) | -7.4 % | <0.0001 | 0.440 ± 0.013 (8) | 0.403 ± 0.005 (14) | -8.4 % | 0.006 |
| 9 months | 0.456 ± 0.003 (12) | 0.416 ± 0.003 (16) | -8.8 % | <0.0001 | 0.467 ± 0.005 (13) | 0.419 ± 0.001 (10) | -10.3 % | <0.0001 |

To determine to what extent specific brain sub-regions are affected, we acquired T2-weighted *in vivo* MRI scans from 4-months old mice (Figure 3-figure supplement 1A). We found significant volume reductions in grey and white matter regions, e.g. in the corpus callosum (-10.2 %), cerebellar grey matter (-7.7 %), cerebellar white matter (arbor vitae, -9 %), hippocampus (-10.7 %), hypothalamus (-4.9 %), striatum (-6.9 %), pons (-9.5 %), medulla (-7.3 %) and cerebral cortex (-8.3%) of *Trappc9* KO mice (Figure 3A, Table 3). When analysed for disproportionately affected brain sub-regions, the hippocampus showed a small, but highly significant decrease in proportional contribution to total brain volume (Figure 3B; WT: 4.51 ± 0.199 %; KO: 4.39 ± 0.168 %, p<0.01), indicating that this brain region is reduced above-average.

**Figure 3.**
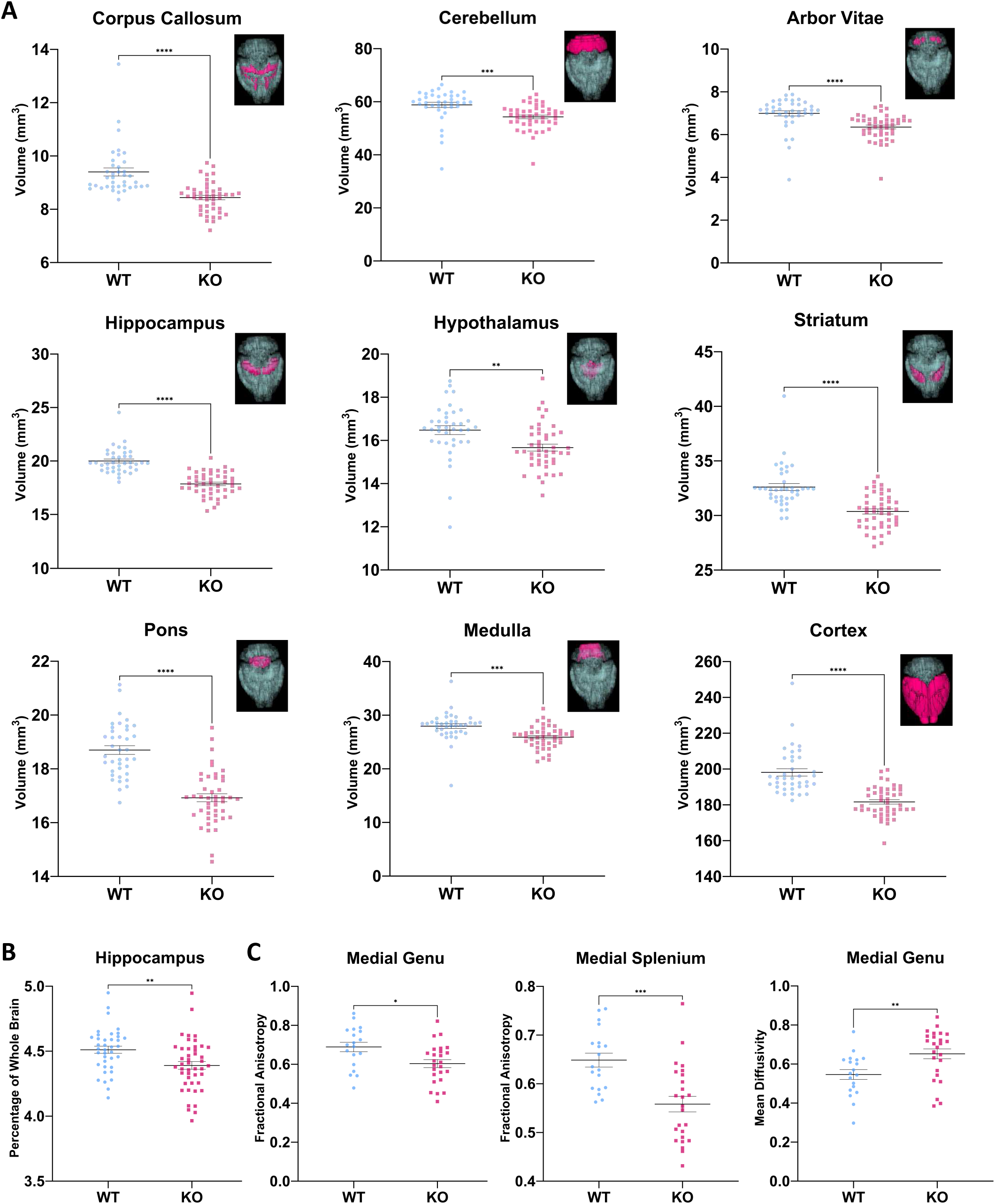
Brain subregion volumes and corpus callosum DTI data. **A)** AIDAmri analysis for brain subregion volumes from T2-weighted *in vivo* MRI scans at age 4 months. Example images for the analysed brain areas are included with each graph, respectively. Table 3. **B)** Hippocampus volume as a proportion of the total brain volume. **C)** Diffusion tensor imaging (DTI) data for corpus callosum sub-regions. Means ± sem; two-way ANOVA with Šídák’s multiple comparison test; *p<0.05, **p<0.01, ***p<0.001, ****p<0.0001.

**Table 3.** Volumes of brain sub-regions in 4-months old WT and *Trappc9* KO mice as measured by T2-weighted *in vivo* MRI and AIDAmri analysis. N = 38 WT and 45 KO mice. Statistical comparisons by two-way ANOVA with Šídák’s multiple comparison test.

| Mean Volume (mm<sup>3</sup>) ± sem |  |  |  |  |
| --- | --- | --- | --- | --- |
| Region | WT | KO | Relative difference to WT | p-value |
| Whole Brain | 443.3 ± 2.50 | 406.6 ± 1.92 | -8.28 % | 0.0001 |
| Cerebral cortex | 198.1 ± 2.07 | 181.7 ± 1.24 | -8.28 % | 0.0001 |
| Corpus callosum | 9.40 ± 0.15 | 8.44 ± 0.08 | -10.21 % | 0.0002 |
| Cerebellar grey matter | 58.83 ± 0.98 | 54.31 ± 0.69 | -7.68 % | 0.0003 |
| Cerebellar arbor vitae | 6.99 ± 0.12 | 6.36 ± 0.09 | -9.01 % | 0.0002 |
| Hippocampus | 20.00 ± 0.19 | 17.86 ± 0.16 | -10.7 % | 0.0002 |
| Hypothalamus | 16.48 ± 0.21 | 15.67 ± 0.16 | -4.92 % | 0.0027 |
| Striatum | 32.61 ± 0.32 | 30.37 ± 0.24 | -6.87 % | 0.0002 |
| Pons | 18.70 ± 0.16 | 16.92 ± 0.15 | -9.52 % | 0.0002 |
| Medulla | 27.96 ± 0.44 | 25.91 ± 0.31 | -7.33 % | 0.0003 |

Using DTI, we determined Fractional Anisotropy (FA) and Mean Diffusivity (MD), which are regarded as measures for white matter integrity and diffusivity in all directions, respectively. We found a significantly lower FA in the medial genu and in the medial splenium region of the corpus callosum (Figure 3C; medial genu: WT: 0.69 ± 0.11 (n = 19), KO: 0.60 ± 0.10 (n = 24), p=0.01; medial splenium: WT: 0.65 ± 0.06 (n = 19), KO: 0.56 ± 0.08 (n = 24), p=0.0002, two-way ANOVA with Šídák’s multiple comparison test). Correspondingly, there was also a significantly higher MD in the medial genu region of the KO corpus callosum (Figure 3C; WT: 0.55 ± 0.11 (n = 19), KO: 0.65 ± 0.13 (n = 24), p=0.005). The changes in FA and MD indicate a more isotropic water diffusion that is less restricted in dimensions by axonal membranes, which allows the conclusion of less white matter organisation or reduced white matter integrity in the KO corpus callosum. We did not detect any differences in FA or MD in the cerebellar white matter.

To follow up on the specific finding of an above-average reduction in hippocampus volumes of *Trappc9* KOs, we investigated the neurogenic niche of the dentate gyrus, which retains NSPCs into adulthood. Using RNAscope *in situ* hybridisation on brain sections of 3-months old mice we found *Trappc9* to be co-expressed with the NSPC marker *Sox2* in cells of the neurogenic subgranular zone, in addition to the very prominent *Trappc9* expression in the neuronal granule cell layer (Figure 4A). Using immunohistochemistry, we quantified Sox2-positive NSPCs within the dentate gyri of 3-months old mice and found a 13 % reduction in their population in *Trappc9* KO samples, especially in the anterior regions of the dentate gyrus (WT: 207.7 ± 7.8 cells per section; KO: 182.0 ± 7.7 cells per section; p<0.05) (Figure 4B). The lower number of NSPCs might, therefore, be a contributing factor for the disproportionately smaller hippocampus volume of KOs.

**Figure 4:**
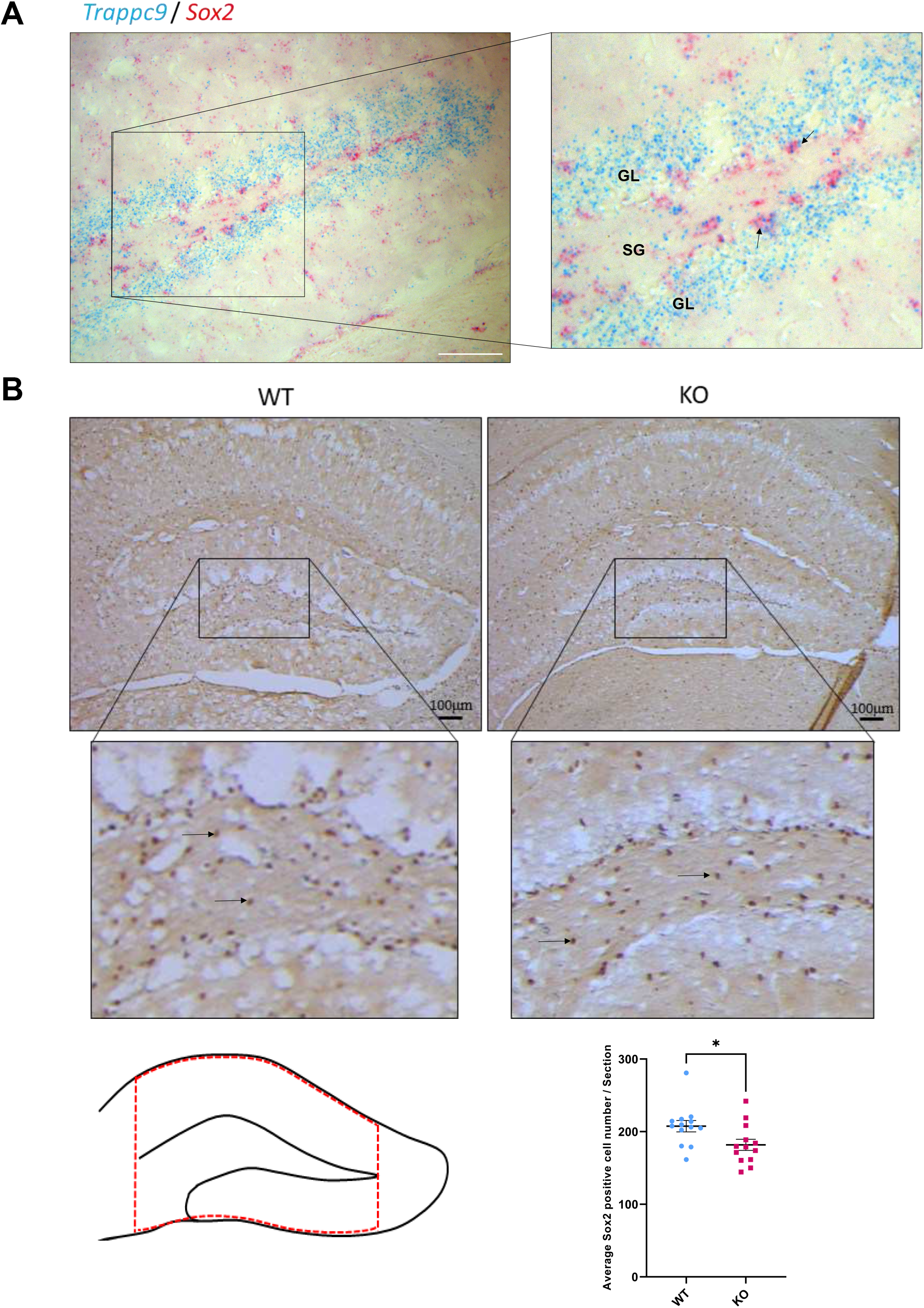
Reduction of Sox2-positive NSPCs in *Trappc9* KO dentate gyrus at age 3 months. **A)** *In situ* hybridisation for *Trappc9* and *Sox2*, which shows co-localisation in the subgranular zone (SGZ) of the dentate gyrus. GL: granular layer. **B)** Immunohistochemistry for Sox2 (arrows) in the hippocampal dentate gyrus of WT and KO. Scheme of the area analysed (delineated by the red dashed line) and quantification of positive cells in this area. Means ± sem. n=3 WTs and 3 KOs (13 averages from brain sections per genotype); independent *t*.test, p: *<0.05. All scale bars: 100 µm.

### Behavioural abnormalities in *Trappc9*-deficient mice include deficits in learning and memory

Since human *TRAPPC9* mutations are associated with severe intellectual disability and to explore whether the microcephaly in KO mice is associated with behavioural deficits, we undertook a series of behavioural tests using the same cohort of mice (at age 4 – 6 months) that underwent *in vivo* MRI. We used the Open Field test to investigate locomotion and anxiety-related behaviour. *Trappc9* KO mice entered the brightly illuminated centre of the open field less often than their WT littermates, and their latency to initially enter the centre was increased (Figure 5A, Table 4). KO mice also spent less time in the centre of the open field. Furthermore, the total distance travelled over the 10 min test time was reduced in the KO mice (Figure 5A, Table 4). The two-way ANOVA indicated that there was no significant interaction between the effects of sex and genotype on any of these data. Simple main effects analysis showed that genotype, but not sex, had a significant effect on the measured parameters. Overall, these data are indicative of increased anxiety and reduced locomotion in the open field context.

**Figure 5.**
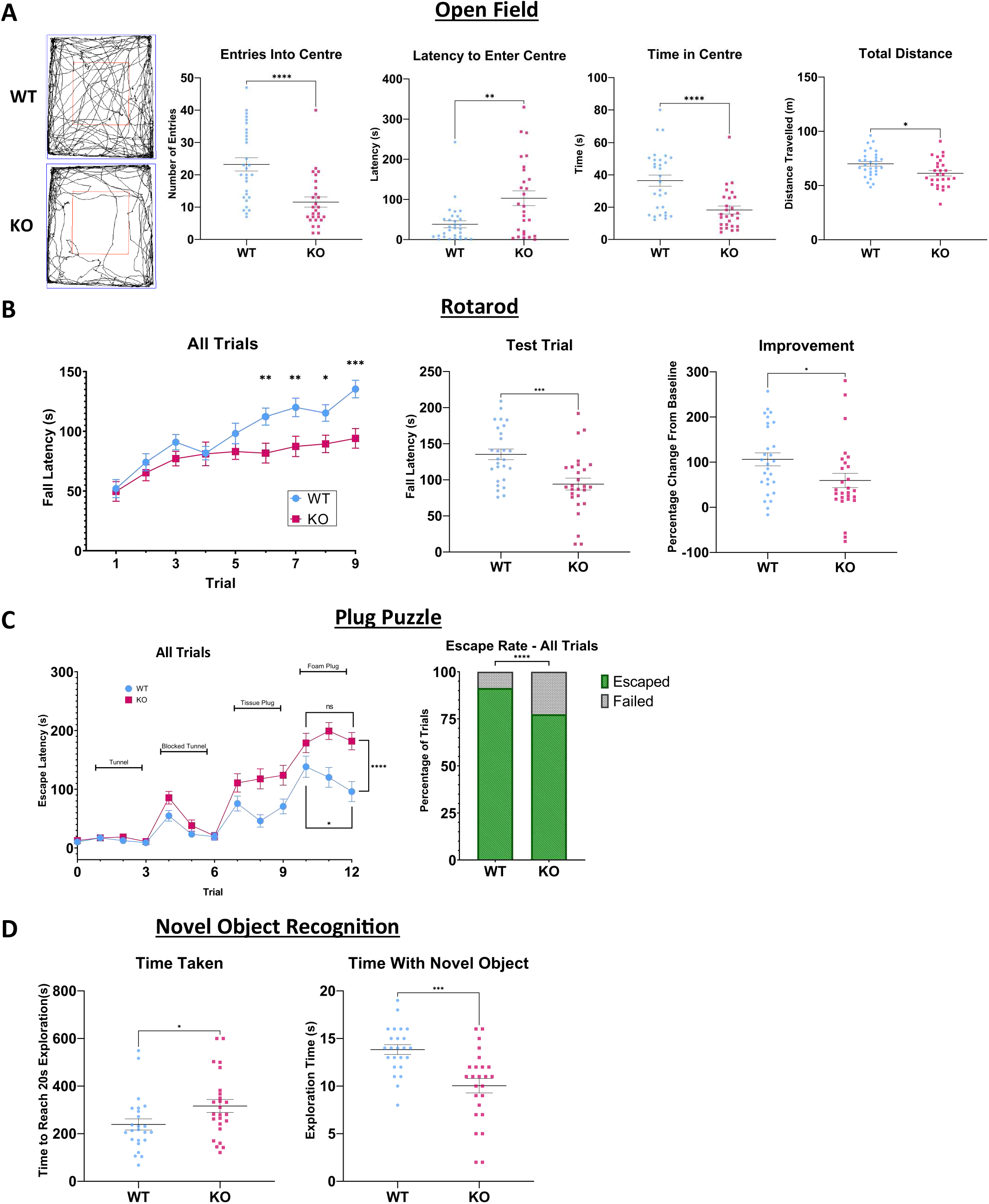
Behavioural tests. **A)** Open Field test. Panels show example tracks and scatterplots for entries into the centre, latency to enter centre, time in centre and total distance moved. **B)** Rotarod test. Panels show fall latency as average performances across all trials, fall latency in the final test trial 9 and improvement from baseline in the test trial 9. **C)** Plug puzzle test. Escape latencies across all trials for WT and KO mice are shown. Differences are indicated for performance between genotypes in final trial 12 and for learning effects within genotypes between trial 10 and 12. The escape / failure rate is shown on the right as a percentage of all trials (Fisher’s Exact test). **D)** Novel object recognition test. The panels show time taken to reach 20 s of active object exploration (both objects combined) and time spent with the novel object (out of the 20 s total object exploration time). Means ± sem. Two-way ANOVA with Šídák’s multiple comparison test; *p<0.05, **p<0.01, ***p<0.001, ****p<0.0001. Table 4.

**Table 4.** Measurements from behavioural tests (means ± sem). Sample size for each test is provided in brackets. Statistical comparison by Two-way ANOVA with Šídák’s multiple comparison test or Fischer’s Exact for the Plug Puzzle – Failure Rate All Trials.

| Behavioural measure<br>(N: WT, KO) | WT | KO | Relative<br>Difference to<br>WT | p-value |
| --- | --- | --- | --- | --- |
| **Open Field (30, 27)** |  |  |  |  |
| Time in Centre (s) | 36.48 ± 3.43 | 18.29 ± 2.48 | -49.86 % | <0.0001 |
| Latency to Enter (s) | 38.26 ± 8.66 | 103.04 ± 18.19 | +169.32 % | 0.0016 |
| Number of Entries | 23.23 ± 2.06 | 11.56 ± 1.58 | -50.24 % | <0.0001 |
| Total Distance (m) | 70.08 ± 2.16 | 61.35 ± 2.48 | -12.46 % | 0.01 |
| **Rotarod (27, 27)** |  |  |  |  |
| Trial 9 Fall Latency (s) | 135.40 ± 7.40 | 94.11 ± 8.24 | -30.49 % | 0.0005 |
| Improvement from Baseline (%) | 106.23 ± 14.27 | 59.51 ± 15.92 | -43.98 % | 0.033 |
| **Plug Puzzle (32, 29)** |  |  |  |  |
| Final Trial Escape Latency (s) | 95.97 ± 17.07 | 181.83 ± 14.72 | +89.47 % | <0.0001 |
| Failure Rate All Trials (%) | 8.65 | 22.55 | +160.69 % | <0.0001 |
| **Novel Object Recognition (24, 25)** |  |  |  |  |
| Time Taken (s) | 238.9 ± 23.60 | 316.6 ± 26.97 | +32.52 % | 0.0359 |
| Time with Novel Object (s) | 13.83 ± 0.506 | 10.04 ± 0.7582 | -27.40 % | 0.0001 |

As a measure of motor coordination and learning, we undertook the accelerating rotarod test. We gave the animals 3 trials / day for 3 days with the trial 9 on day 3 acting as the final indicative test. Although fall latency was not different in the first trial on day 1, KO mice fell from the beam earlier than WTs in the final test trial 9 (Figure 5B, Table 4). To calculate improvement in the balancing task, we averaged the first day’s trials 1 – 3 as a baseline and the percentage change from this in the final trial 9. The overall improvement from the baseline score was lower in the KOs than in WT littermates (Figure 5B, Table 4). A two-way ANOVA revealed that there was no significant interaction between the effects of sex and genotype on rotarod fall latency or improvement. Simple main effects analysis showed that genotype had a statistically significant effect on both parameters while sex did not. Overall, the data from the rotarod tests show no basic motor coordination fault in *Trappc9* KO mice, but a reduced capacity in learning to improve motor coordination.

As a test for cognitive problem solving and memory, we carried out the Plug Puzzle test (O’Connor et al., 2014), which consists of a brightly lit open field and a dark escape box separated by a doorway, which is plugged with increasingly difficult obstacles that the mouse must remove. KO mice took longer to escape in the final probe trial of the Plug Puzzle test than their WT littermates (Figure 5C, Table 4). While WT mice showed a learning effect over the foam plug trials 10 – 12, KO littermates failed to improve over the three repeats (Figure 5C; WT trial 10: 138.3 ± 18.01 s, trial 12: 95.97 ± 17.07 s, p=0.011; KO trial 10: 178.7 ± 16.54 s, trial 12: 181.8 ± 14.72 s, p=0.82 n.s.; mean ± sem; paired *t*.test). A two-way ANOVA revealed that there was no significant interaction between the effects of sex and genotype in the final trial of the Plug Puzzle. Simple main effects analysis showed that genotype did have a significant effect on escape latency (Table 4), but sex did not. Considering all trials, KO mice were also three times more likely to fail a trial (Figure 5C, Table 4). Thus, the results of the Plug Puzzle test indicate reduced cognitive problem-solving capability and memory in *Trappc9* KO mice.

To examine non-spatial memory and exploratory behaviour, we carried out the Novel Object Recognition test (Leger et al., 2013), which is based on the assumption that a mouse that memorises a familiar object will explore a novel object more. After initial habituation and object familiarisation stages, we presented the mice with a novel object alongside the familiar one during the test trial. KO mice took longer than WTs to reach 20 seconds of total object exploration (Figure 5D, Table 4). Out of the 20 seconds total object exploration, KOs spent less time interacting with the novel object (Figure 5D, Table 4). A two-way ANOVA revealed that there was no significant interaction between the effects of sex and genotype in the time taken for exploration or in time spent with the novel object. Simple main effects analysis showed that genotype did have a significant effect on time taken and on time spent with the novel object while sex did not (Table 4). These data show that *Trappc9*-deficient mice performed worse in object recognition memory indicating a lack of curiosity (Leger et al., 2013).

### Lipid droplet homeostasis is perturbed in primary Trappc9-deficient neuron cultures

*Trappc9* deficiency in immortalised cell lines and patient fibroblasts results in a disturbed LD homeostasis and impaired lipolysis, which is due to disruption of TrappII-mediated guanine nucleotide exchange and activation of the LD-specific protein Rab18 (Li et al., 2017). To investigate whether these cellular phenotypes can be observed in disease-relevant neuronal cells, we analysed LDs in cultured hippocampal neurons from *Trappc9* KO mice after 6 and 12 hrs incubation with OA. Although the total LD volume per cell was not different between genotypes after 6 hrs of OA (WT: 61.76 µm^3^ ± 8.70, KO: 83.19 µm^3^ ± 11.74, p=0.14, independent *t*.test), it became significantly larger in KO neurons at 12 hrs (WT: 77.78 µm^3^ ± 8.53, KO: 129.7 µm^3^ ± 12.52, p=0.001, independent *t*.test), due to a strong increase between the two time points (p=0.009) (Figure 6A, B). Analysis of the sizes of individual LDs indicated that LD volumes were larger in KO than in WT neurons at 6 hours (WT: median 1.67 μm^3^, IQR 0.62 – 3.8; KO: median 2.68 μm^3^, IQR 1.04 – 6.06; p<0.0001, Mann-Whitney U-test) (Figure 6C). At the 12 hrs time point this difference was lost, although a trend to increased individual LD sizes remained in the KO neurons (WT: median 2.82 μm^3^, IQR 1.07 – 5.80; KO: median 2.99 μm^3^, IQR 0.92 – 7.47; p=0.21, Mann-Whitney U-test). The volumes of individual LDs expanded significantly from 6 to 12 hrs in WT neurons (p<0.0001) (Figure 6C), while in KO neurons only a trend towards a size increase could be observed between these two time points. These results indicate that individual LDs grow more quickly in the KO hippocampal neurons compared to WT, eventually resulting in a larger total LD volume per cell after 12 hrs of OA exposure.

**Figure 6.**
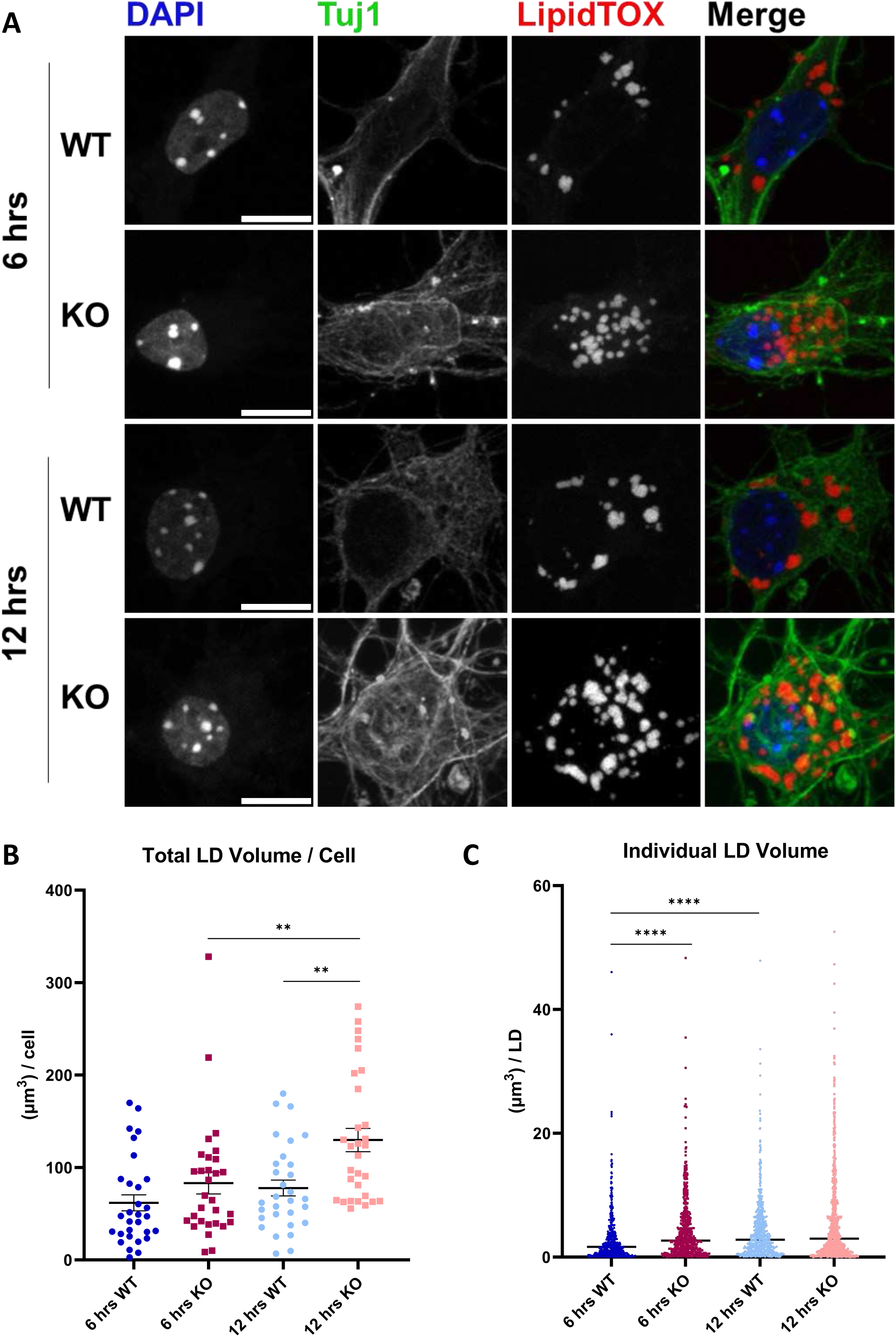
Abnormal LD accumulation in KO hippocampal neurons. **A)** Representative images of neurons (Tuj1) after 6 and 12 hrs of OA incubation (LDs stained with LipidTOX). Scale bar: 10 µm. **B)** Quantification of total LD volume per cell. Each dot represents a single cell; means ± sem; total n = 120 cells from 3 animals (∼10 cells / animal / group), independent *t*.test. **C)** Quantification of individual LD volumes. Each dot represents a single LD; median ± IQ range; Mann-Whitney U-test (total n = 2376 LDs). **p<0.01; ****p<0.0001.

To further investigate LD homeostasis, we analysed the association of Plin2/Adrp with LD surfaces, since Plin2 is involved in the regulation of lipolysis and lipophagy (Sztalryd & Brasaemle, 2017) and has been found to co-localise with Trappc9 at LD surfaces (Li et al., 2017). Both, Plin2 and Trappc9, also interact with Rab18 (Deng et al., 2021). Furthermore, Plin2 is the most highly expressed perilipin in NSPCs of the dentate gyrus and subventricular zone (Inloes et al., 2018; Ramosaj et al., 2021). In WT hippocampal neurons, we found a prominent localisation of Plin2 around LDs, especially at the earlier time point of 6 hrs after OA supplementation, while this association was much reduced in *Trappc9* KO neurons at both time points (Figure 7A). Quantification showed that the vast majority of LDs were Plin2-positive at 6 hrs of OA in WT and KO neurons, with a trend towards a larger percentage in KOs (WT: 77.0 %; KO: 81.6 %; p=0.05, Chi-square test) (Figure 7B). At 12 hrs of OA, the percentage of Plin2-positive LDs decreased slightly, but significantly, in WT neurons while a much stronger reduction was observed in KO neurons, which resulted in a significant difference between genotypes (WT: 70.5 %; KO: 62.0 %; p<0.001, Chi-square test) (Figure 7B). To characterise the Plin2 phenotype in more detail, we determined the proportion of the LD surface area associated with Plin2 staining in the Plin2-positive LD groups. The portion of LD surface area coated by Plin2 varied considerably in all groups analysed, and Plin2 localisation around LDs decreased significantly between 6 and 12 hrs of OA in both WT and KO neurons (Figure 7C). However, compared to WT, *Trappc9* KO neurons showed overall significantly less association of Plin2 with LD surfaces at both time points of OA supplementation (Figure 7C).

**Figure 7.**
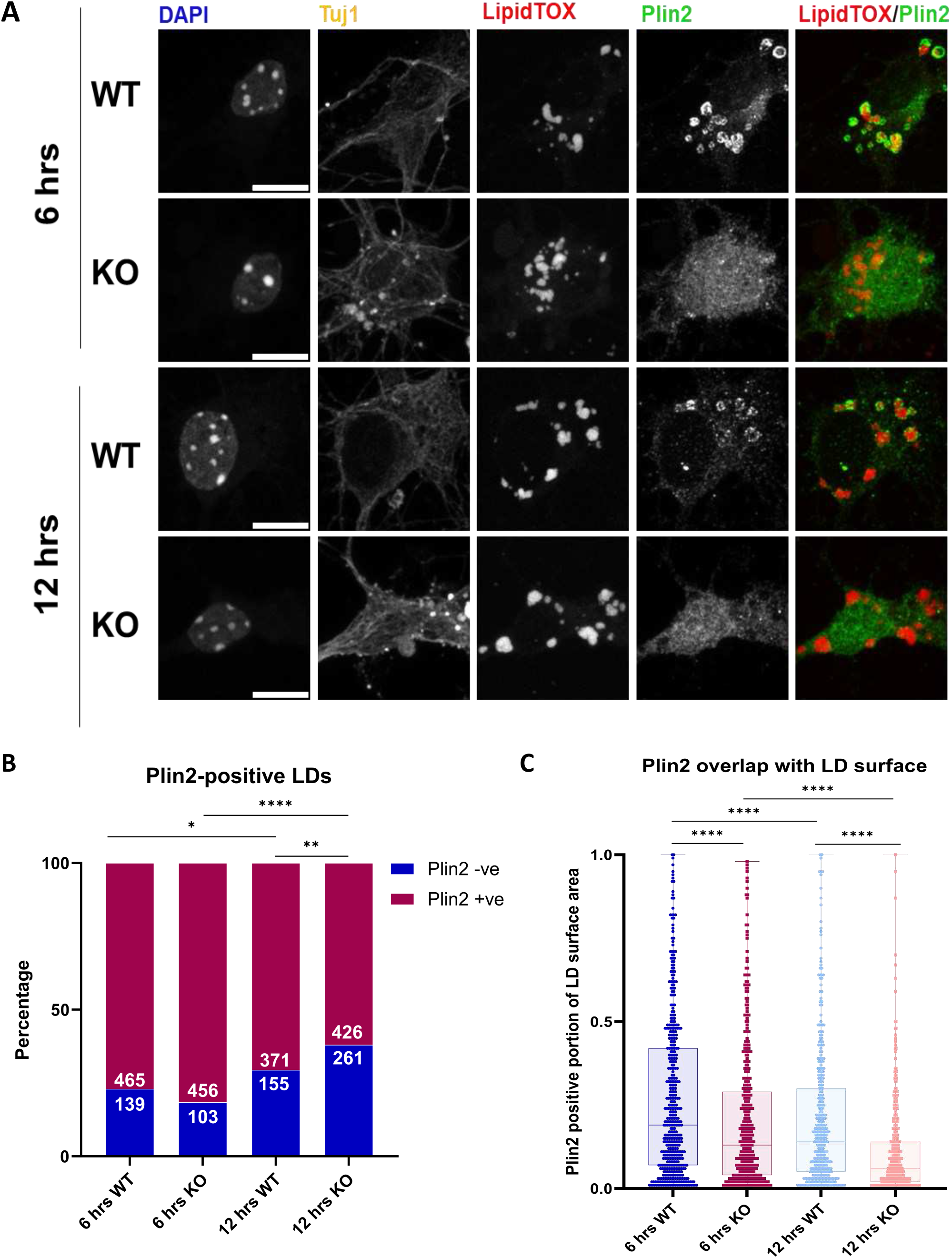
Reduced association of Plin2 with LDs in *Trappc9* KO hippocampal neurons. **A)** Representative images of co-staining with neutral lipid dye LipidTOX and lipid droplet protein Plin2 after 6 and 12 hrs of Oleic Acid supplementation. Scale bar: 10 µm. **B)** Percentage of Plin2-positive and negative LDs (Chi-square test). **C)** Quantification of the portion of LD surface area coated by Plin2 (only Plin2-positive LDs were analysed). Each dot represents a single LD (total n = 1718 LDs). Median ± IQR; Mann-Whitney U-test. *p<0.05; **p<0.01; ****p<0.0001.

Our data of increased lipid storage and reduced Plin2-coating of LDs in *Trappc9* KO neurons indicate that the regulation of LDs is impaired, which might impact on lipid and/or fatty acid metabolism and lipotoxicity in neural cells (Ioannou, Jackson, et al., 2019; Kandel et al., 2022; Ralhan et al., 2021; Ramosaj et al., 2021).

### *Trappc9*-deficient mice develop an obesity phenotype that is more severe in females than in males

Obesity is one of the symptoms frequently reported in *TRAPPC9*-deficient human patients (Aslanger et al., 2022; Bolat et al., 2022; Kramer et al., 2020). To assess this phenotype in mice, we monitored their body weight across age. *Trappc9* KO mice showed no difference in body weight on the day of birth or at one month of age (Figure 8A, B). A significantly higher body weight was first observed in female KOs at two months, and this difference increased steadily during adult stages (Figure 8C). By contrast, overweight in male KOs became apparent at seven months only (Figure 8D). The body weight increase was significantly higher in female (+29 %) than in male (+ 9 %) KO mice when normalised to the average of their same-sex WT littermates at nine months of age (Figure 8E). While there was no difference in blood glucose levels of *ad libitum* fed mice (Figure 8F), plasma leptin was elevated in female KOs (Figure 8G). Furthermore, histological sections of white and brown adipose tissues showed increased adipocyte and lipid droplet sizes in KO mice of both genders (Figure 8-figure supplement 1). Taken together, these results indicate an obesity phenotype in *Trappc9*-deficient mice, which is significantly more pronounced in females and develops after the onset of microcephaly.

**Figure 8.**
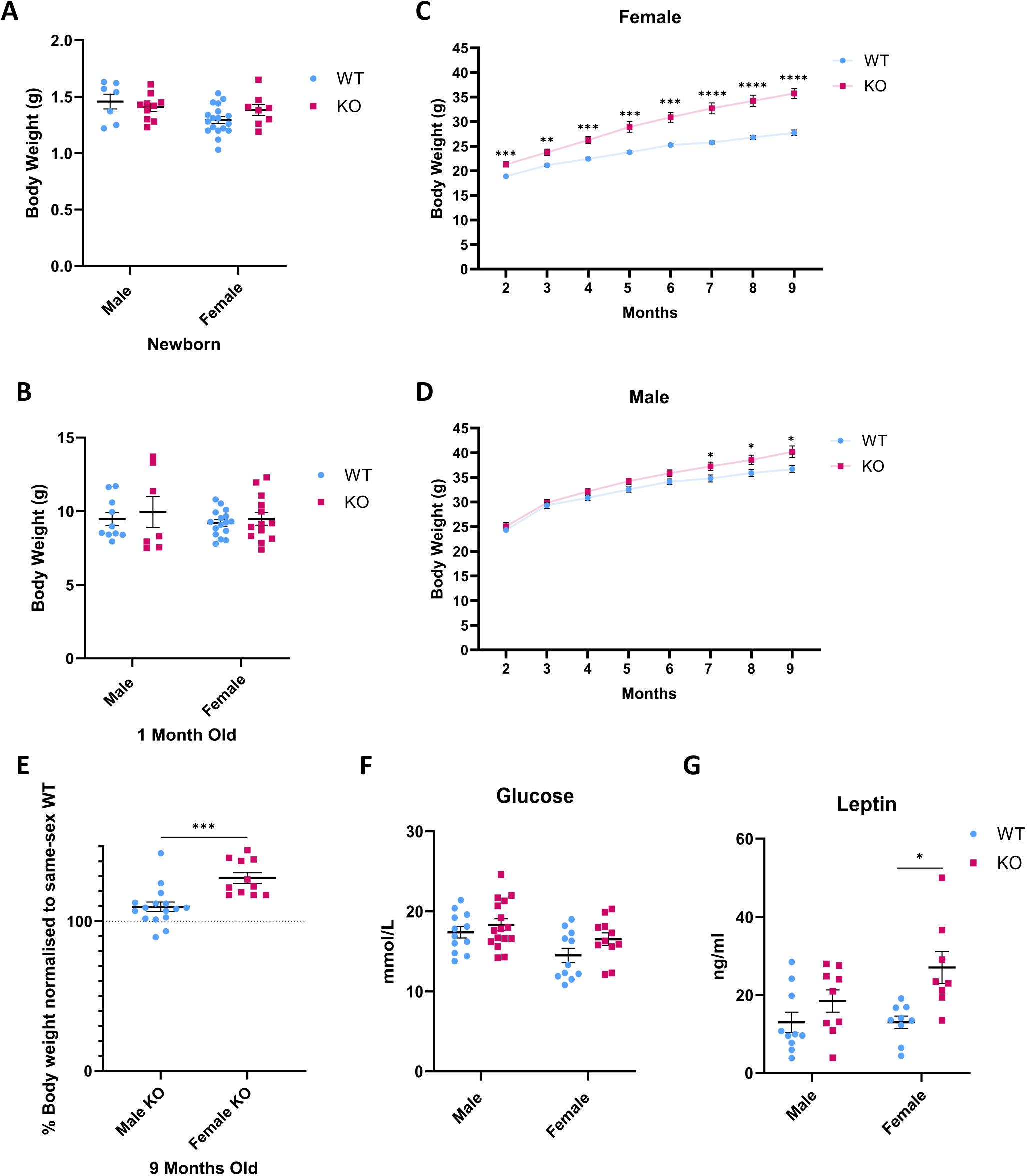
Obesity phenotype is more severe in *Trappc9* KO females. **A)** and **B)** Body weights of newborn and one month-old mice. **C)** and **D)** Body weights of male and female mice at adult stages. **E)** Overweight is significantly higher in female KOs than in male KOs. Data are normalised to the average of same-sex WT littermates. **F)** Plasma blood glucose and **G)** Plasma Leptin levels of 9-months old mice. Female KOs have increased leptin levels. Means ± sem. Independent *t-test*. *p<0.05, **p<0.01, ***p<0.001, ****p<0.0001.

## Discussion

In this study, we demonstrate by using MRI that the microcephaly of *Trappc9* KO mice has a postnatal onset and is clearly established at weaning age. These findings are in line with *TRAPPC9* patient data, which show microcephaly within the first year of life (Amin et al., 2022; Aslanger et al., 2022; Ben Ayed et al., 2021; Bolat et al., 2022; Hnoonual et al., 2019; Koifman et al., 2010; Penon-Portmann et al., 2023; Radenkovic et al., 2022) as well as data from other recently published *Trappc9* KO mouse studies, which reported differences at postnatal days 7, 15 and 20, but not at birth (Hu et al., 2023; Ke et al., 2020). Monogenic disorders causing postnatal-onset microcephaly are less common than those causing primary microcephaly, which are mostly due to cell proliferation defects during embryogenesis. However, it has been noted that postnatal microcephaly, white matter defects and intellectual disability often occur when genes related to Golgi apparatus functions are mutated, for which the term ‘Golgipathies’ was proposed (Rasika et al., 2018). This category includes many small Rab GTPases as well as their GEFs and GTPase-activating proteins (GAPs), which are involved in the regulation of intracellular membrane compartments and trafficking processes that become especially important during postnatal neuronal differentiation and maturation (Rasika et al., 2018). Since the Trappc9-containing TrappII complex acts as a GEF for the endosome-recycling and Golgi-associated Rab11 (Jenkins et al., 2020; Ke et al., 2020), *Trappc9*-related postnatal microcephaly can be considered a ‘Golgipathy’. This suggestion is further supported by findings of endoplasmic reticulum to Golgi transport defects and Golgi fragmentation in cells deficient for the TrappII-specific subunit TRAPPC10 or the core subunits TRAPPC6B, TRAPPC4 and TRAPPC2L, which all lead to disruption of a normal TrappII complex and similar neurodevelopmental disorders (Al-Deri et al., 2021; Almousa et al., 2023; Rawlins et al., 2022; Van Bergen et al., 2020). These neurodevelopmental disorders constitute a subset within the spectrum of ‘Trappopathies’ caused by mutations in genes for Trapp subunits (Sacher et al., 2019).

Our detailed analysis for volume differences in specific brain sub-regions at the adult stage via *in vivo* MRI showed a similar reduction in most grey and white matter regions, apart from the hippocampus, which was disproportionately more severely affected. These findings differ partly from histology-based brain morphometry data, which describe decreases in only a limited set of *Trappc9* KO brain regions (Ke et al., 2020; Liang et al., 2020). While the corpus callosum was consistently found to be reduced in all *Trappc9* mouse studies (Hu et al., 2023; Ke et al., 2020; Liang et al., 2020), our data contrast with the volume increase described for the striatum by Ke et al. (2020) (Ke et al., 2020). The discrepancies in the analysis of brain sub-regions remain to be resolved, but can most likely be attributed to differences in methodology and sample size. Our *in vivo* MRI analysis of the unperturbed brain allowed us to process a large number of samples without tissue sectioning and histological processing. The corpus callosum is not the only white matter fibre tract affected, as we found the arbor vitae of the cerebellum to be reduced as well, which is in line with findings of smaller nerve bundles in the striatum and reduced white matter in the spinal cord described by Hu et al. (2023) (Hu et al., 2023). Our *in situ* hybridisation analysis indicates that *Trappc9* is mainly expressed in neuronal cell areas, and the single cell analysis of the Allen Brain Cell Atlas (https://portal.brain-map.org/atlases-and-data/rnaseq) (Yao et al., 2021) shows highest expression levels of *Trappc9* in in a wide range of cortical and hippocampal neurons with much lower levels found in glial cells and oligodendrocytes. This is consistent with immunohistochemistry data by Ke et al. (2020) (Ke et al., 2020) and indicates that the white matter reductions are primarily due to axonal deficiencies with decreases in myelin being a secondary effect (Hu et al., 2023). Furthermore, a single cell analysis of newborn *Trappc9* KO cerebral cortex did not detect any major changes in neural cell type composition, but did identify gene expression changes in pathways of neuritogenesis, synaptogenesis, vesicle trafficking and intracellular membrane compartments (Hu et al., 2023). In this context, our white matter corpus callosum DTI data of reduced fractional anisotropy and increased mean diffusivity might be due to decreased axonal organisation or alignment, although we cannot currently rule out reduced myelination as a contributing factor.

We found the hippocampus to be the only region that showed a disproportionately greater volume reduction relative to the whole KO brain. Furthermore, we show that *Trappc9* is highly expressed not only in granule neurons of the hippocampus, but also in adult NSPCs of the sub-granular zone of the dentate gyrus, and that the number of Sox2-positive NSPCs in this region is reduced in 3-months old KOs. Since the hippocampus is one of only two brain regions containing adult NSPCs (Denoth-Lippuner & Jessberger, 2021; Goncalves et al., 2016), its disproportionate volume reduction might be due to a deficiency in adult NSPC proliferation, survival or differentiation in addition to defects in mature neurons that also occur in other KO brain regions. Although these findings require further investigation, they are supported by similar observations of reduced NSPCs in 3-week old *Trappc9*-null mice (Usman et al., 2022).

In addition to Rab11, Rab18 was shown to be another substrate for GEF activity by the TrappII complex in the context of LD regulation, whereby Trappc9-deficient patient fibroblasts as well as cell lines lacking either Trappc9, Trappc10 or both displayed increased LDs sizes (Li et al., 2017). We investigated this phenotype in disease-relevant primary hippocampal KO neurons, since neurons are sensitive to lipotoxicity and normally do not contain significant amounts of LDs, but on the other hand require lipids for membrane formation during periods of neurite outgrowth and can form LDs when lipid metabolism is disturbed (Chung et al., 2023; Inloes et al., 2014; Ioannou, Jackson, et al., 2019; Ralhan et al., 2021). Our data support a role of Trappc9 in LD regulation, as KO neurons accumulated a larger total LD volume per cell and individual LDs were larger during early stages of OA exposure, indicating a quicker LD growth. Furthermore, we found that the portion of LD surface areas coated by Plin2/Adrp, one of the major LD-associated proteins (Sztalryd & Brasaemle, 2017), was much reduced in *Trappc9* KO neurons while the percentage of LDs that lacked Plin2 completely was increased. Trappc9 co-localises with Plin2/Adrp during early stages of LD formation and both, the TrappII complex and Plin2, bind to Rab18 and facilitate its recruitment onto LDs (Deng et al., 2021; Li et al., 2017). These findings suggest an interaction between the three proteins on LD surfaces, which might be of functional importance in the regulation of lipid homeostasis in neurons. It is noteworthy in this context that mutations in *RAB18* lead to Warburg Micro syndrome, a neurodevelopmental disorder with some similarities to *TRAPPC9* deficiency, including postnatal microcephaly, intellectual disability and enlarged LDs (Bekbulat et al., 2020; Bem et al., 2011; Carpanini et al., 2014; Xu et al., 2018). Neuronal LD phenotypes similar to those described here for Trappc9 deficiency have also been identified in two other neurobiological disorders, Troyer syndrome and SPG54, which are regarded as specific sub-forms of hereditary spastic paraplegia (Chung et al., 2023; Inloes et al., 2014). The associated genes *SPARTIN* and *DDHD2* function in autophagy of LDs and triglyceride hydrolysis, respectively, while the precise molecular physiology of Trappc9 and the TrappII complex in LD formation and/or degradation remains to be elucidated. Apart from neurons, LDs also have an important role in NSPCs. LD abundance influences their states of quiescence, proliferation or differentiation, and NSPCs of the sub-ventricular and hippocampal sub-granular zones express Plin2 (Ramosaj et al., 2021). Furthermore, NSPCs express the nuclear receptor TLX/NR2E1, which specifically binds Oleic Acid and regulates a set of cell cycle and neurogenesis genes (Kandel et al., 2022). OA application into the dentate gyrus stimulates NSPC proliferation and neurogenesis (Kandel et al., 2022). It is therefore possible that impaired LD homeostasis in *Trappc9* KO mice also affects NSPCs and in this way contributes to the lower number of Sox2-positive NSPCs and disproportionately stronger reduction in hippocampus volume discussed above.

We show that the brain abnormalities identified in *Trappc9* KO mice also result in behavioural deficits related to anxiety, cognition, learning and memory, which might reflect some the intellectual disability symptoms of human patients (Amin et al., 2022; Aslanger et al., 2022; Ben Ayed et al., 2021; Bolat et al., 2022; Hnoonual et al., 2019; Koifman et al., 2010; Kramer et al., 2020; Mir et al., 2009; Mochida et al., 2009; Penon-Portmann et al., 2023; Philippe et al., 2009; Radenkovic et al., 2022). Our open field test data demonstrate an overall reduced locomotor activity of *Trappc9* KO mice, which is consistent with recent findings from other *Trappc9* mouse lines (Hu et al., 2023; Ke et al., 2020; Liang et al., 2020). Furthermore, the KO mice showed increased anxiety to explore the centre of the open field, which was also observed by Ke et al (2020) (Ke et al., 2020), but not by Liang et al (2020) (Liang et al., 2020). This discrepancy can most likely be attributed to experimental design, since in our study, as well as in Ke et al., all behavioural tests were carried out during the wakeful active period of the mice (dark phase of the day), while Liang et al tested during the restive light phase. The rotarod test indicated no difference in fall latency in the first trials. But in contrast to WTs, KO mice only showed a minor improvement in their motor coordination over repeated trials, resulting in increasing performance gaps between the genotypes. While basic initial motor coordination appears to be normal, these data can be interpreted as a limited capacity of the KO mice to learn how to improve their motor coordination. The novel object recognition test (Leger et al., 2013) indicated that *Trappc9*-deficient mice took longer to reach a specified object exploration time and spent less time with the novel object. Similar observations were made in other *Trappc9* mouse lines (Hu et al., 2023; Ke et al., 2020). Overall, these findings confirm a reduced exploratory activity and impaired object memory in the mutant mice. The plug puzzle test, which investigates problem solving and memory abilities (O’Connor et al., 2014), also showed a performance deficit of the KO mice, which took longer to remove the plug and more often failed the task completely at the most difficult stage. In contrast to WTs, they also did not learn over the final three test trials. Taking into account additional behavioural tests undertaken with other *Trappc9* lines, including the Morris water maze, the Barnes maze and social learning tests (Hu et al., 2023; Ke et al., 2020; Liang et al., 2020), it can be concluded that lack of Trappc9 in mice leads to cognitive, memory and learning impairments.

Apart from neurobiological phenotypes, *Trappc9* KO mice develop obesity, which is significantly more severe in females than in males. Increased body weight becomes evident after weaning and is noticeable earlier in females then in males. These findings are consistent between different *Trappc9* mouse lines (Ke et al., 2020; Liang et al., 2020). Adipocytes in brown and white adipose tissue show an increased size and larger lipid droplets, but leptin levels are significantly increased only in female KOs. Similar sex-dependent differences in obesity phenotype were described for *Trappc10* KO mice (Rawlins et al., 2022), which suggests that a dysfunction of the TrappII complex through mutation of either of its two specific subunits, c9 or c10, underlies such shared phenotypes. A more detailed analysis of metabolism in the *Trappc9* KO mice identified hyperinsulinemia, glucose intolerance and increased plasma lipid levels (Liang et al., 2020). It remains unclear whether the obesity phenotype is due to an adipose tissue autonomous function of Trappc9 (Usman et al., 2022), possibly related to LD regulation, or whether impaired brain control of food intake and energy expenditure play a role as well. The latter view is supported by our finding of prominent *Trappc9* expression in hypothalamic paraventricular and arcuate nuclei, which are major centres for the central regulation of energy homeostasis (Bruning & Fenselau, 2023). Furthermore, *Trappc9* is genomically imprinted specifically in the murine brain with a preferential expression bias (∼70 %) from the maternally inherited allele (Claxton et al., 2022; Liang et al., 2020). Accordingly, it has been shown that the phenotypes of *Trappc9* heterozygotes differ depending on the parental inheritance of the mutation. Heterozygotes carrying the mutation on the maternal allele (m-/p+) are almost as severely affected as homozygous KOs, while m+/p- mice are not significantly different from WT in most aspects, including obesity (Liang et al., 2020). Since *Trappc9* shows equal biallelic expression in peripheral tissues and m+/p- mice are not overweight, the obesity phenotype of m-/p+ mice suggests that it is at least partly due to a loss of function in the brain. Our data as well as Ke et al. (2020) (Ke et al., 2020) also indicate that the brain phenotype (microcephaly) develops earlier than the body weight increase, which might therefore be a consequence of the former. Interestingly, the rare metabolic analysis of a human *TRAPPC9* patient indicated hyperphagia as the underlying cause of obesity (Liang et al., 2020). Conditional, tissue-specific *Trappc9* deletions will be required to clarify this phenotype.

Apart from mammalian-specific *Trappc9* KO phenotypes, there are some similarities to mutants in other species. Deletion of the *Drosophila* ortholog *brunelleschi* (*bru*) causes failures in male meiotic cytokinesis due to defects in cleavage furrow ingression in spermatocytes, a process that involves Rab11 (Riedel et al., 2018; Robinett et al., 2009). Male infertility has also been observed in homozygous *Trappc9* KO mice (Hu et al., 2023; Ke et al., 2020). Cytokinesis defects during mitotic cell division have been described for ortholog mutants in the fission yeast *Schizosaccharomyces pombe* and in *Arabidopsis thaliana*, whereby the transport and deposition of cargo materials at the newly forming cell membranes or cell plates, respectively, is impaired (Rybak et al., 2014; Wang et al., 2016). Disruption of the *Saccharomyces cerevisiae* ortholog *Trs120* leads to endosome recycling defects (Cai et al., 2005), which has been confirmed in *Trappc9*-deficient neurons (Ke et al., 2020). Taken together, these findings indicate that Trappc9 and the TrappII complex are involved in multiple functions related to intracellular membrane compartments, some of which are essential and non-redundant in a cell type-specific way. It seems likely that the diverse functions depend on interactions with different Rab and/or other membrane-associated proteins, which remain to be explored.

## Materials and Methods

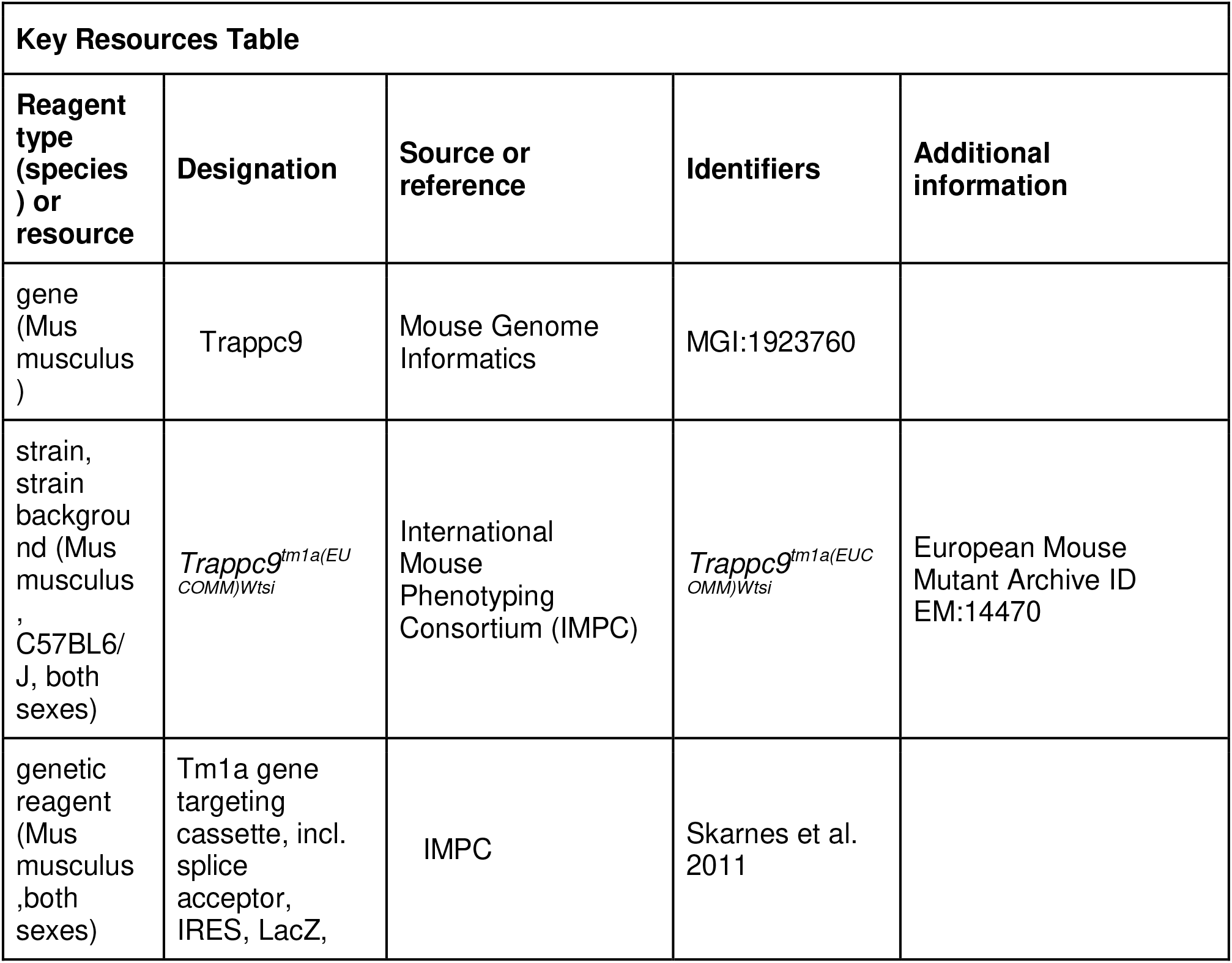
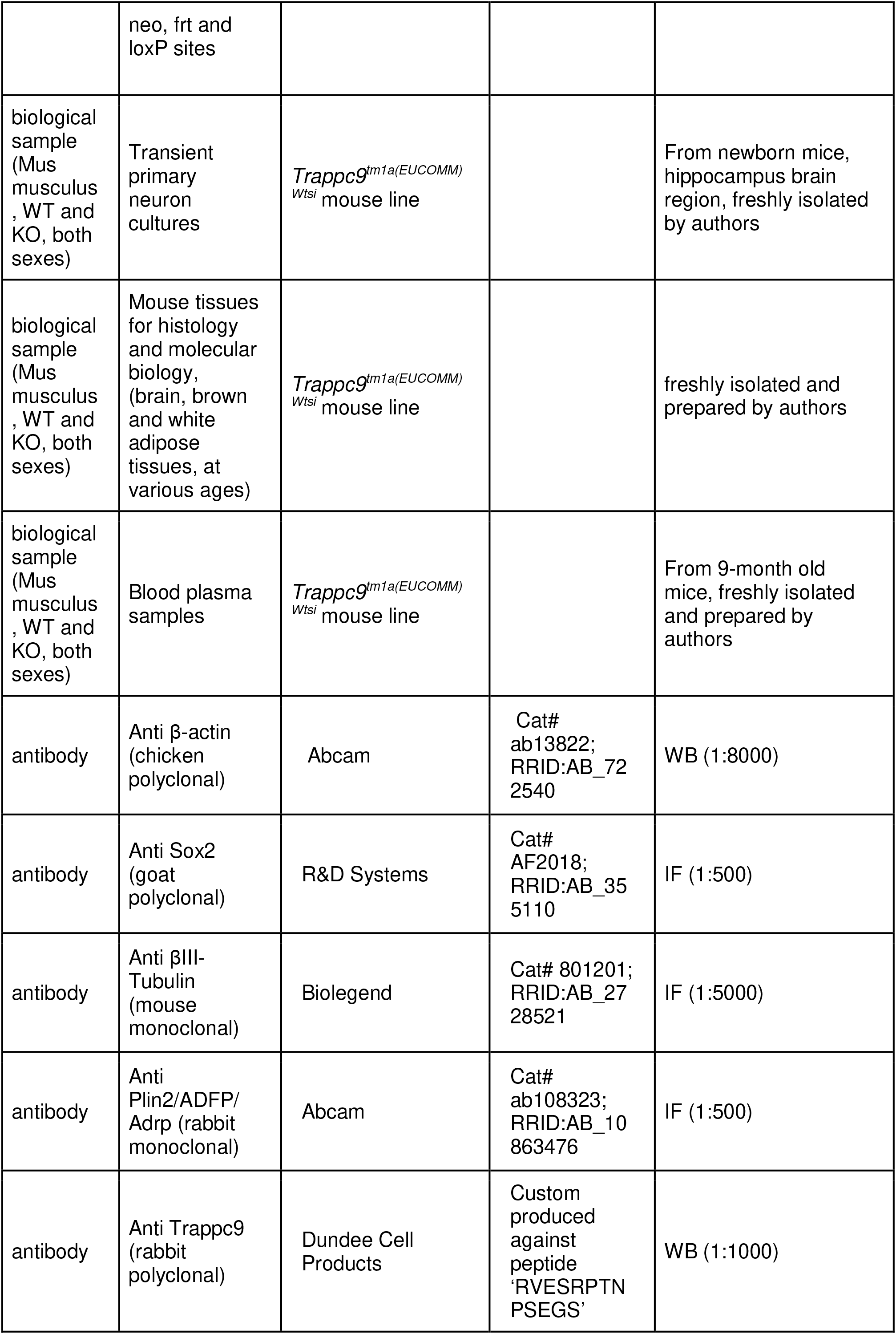
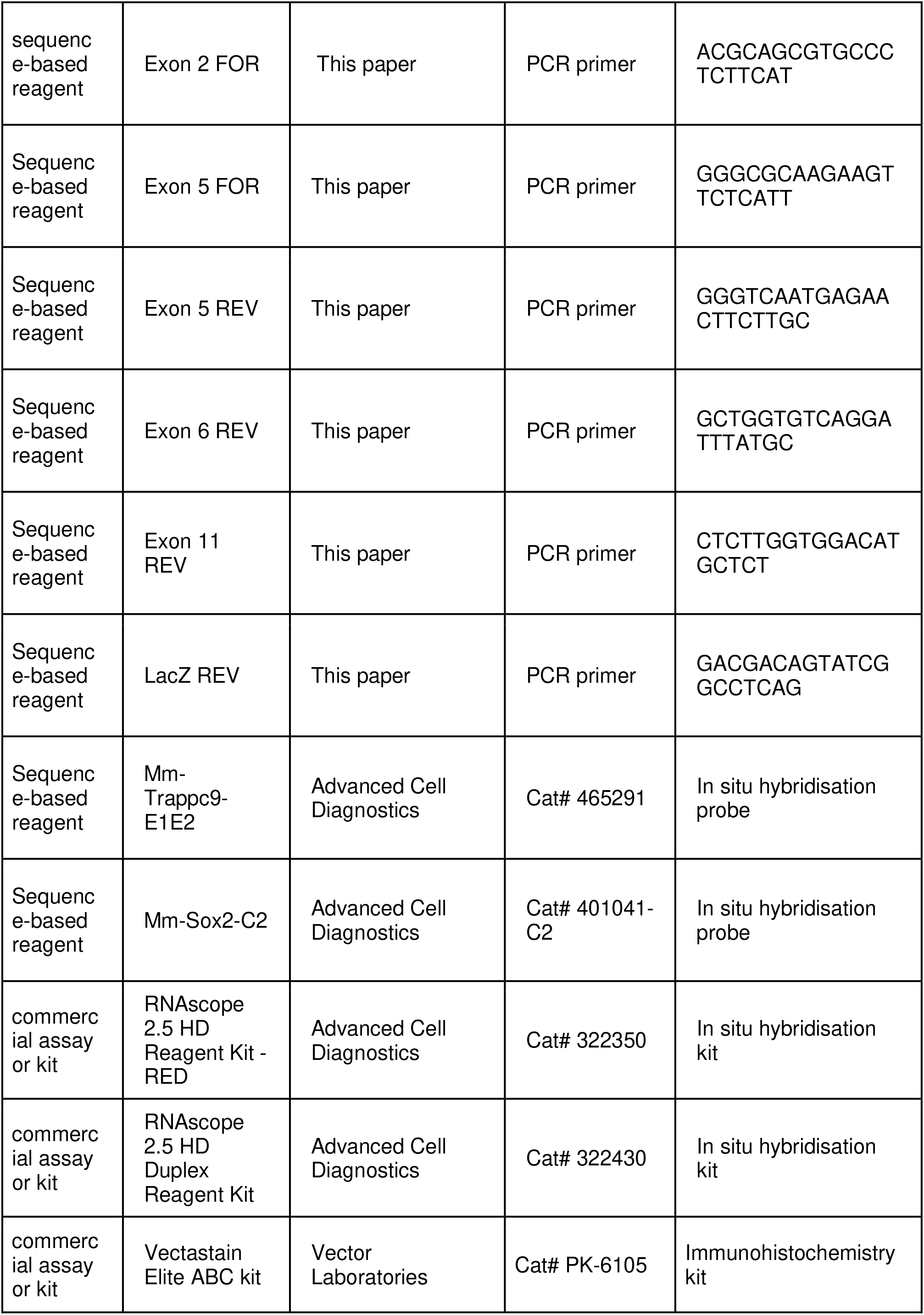
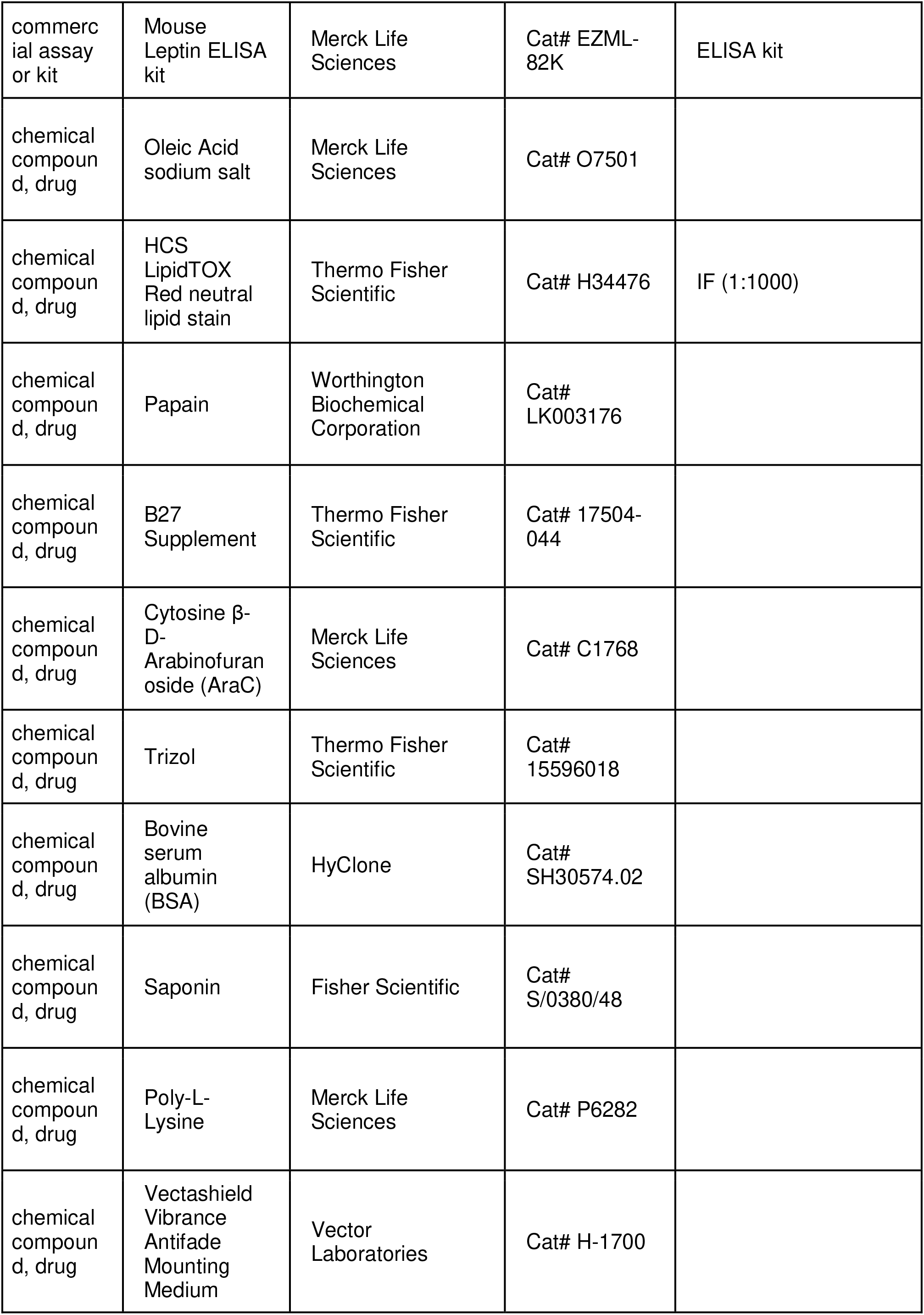
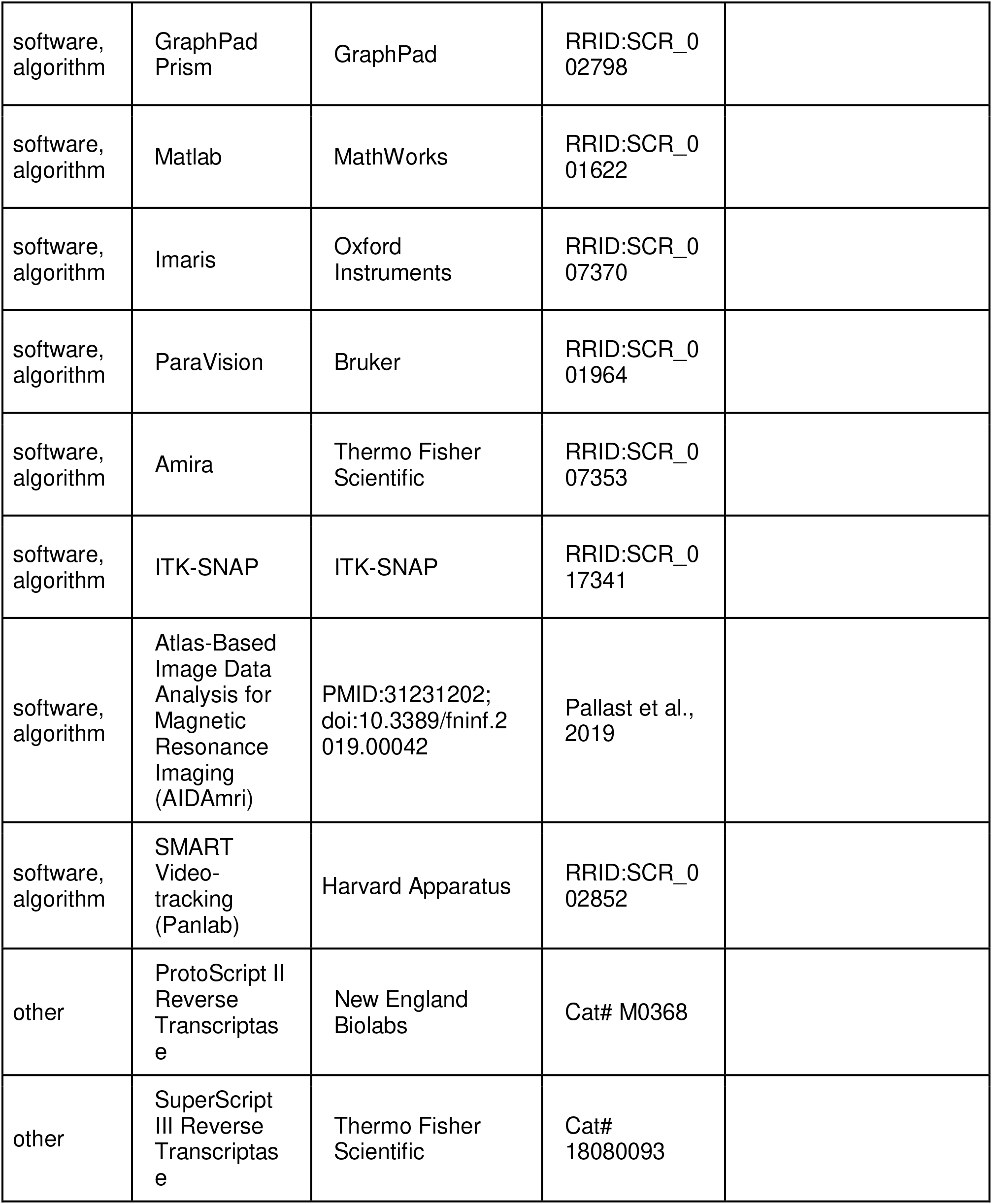

### Animals

The *Trappc9^tm1a(EUCOMM)Wtsi^* knock-out first mouse line (Skarnes et al., 2011) was generated by the International Mouse Phenotyping Consortium (IMPC; www.mousephenotype.org) and imported from the Wellcome Trust Sanger Institute, Cambridge, UK. We maintained the mice at the University of Liverpool Biomedical Services Unit and housed them as same-sex littermate groups in individually ventilated cages with Lignocel Select as the substrate and Z-Nest as the paper-based nesting material. On rare occasions, when no same-sex littermate was available, we housed mice singly. All mice are provided with a balcony, dome home and handling tunnel as enrichment. They were kept under a 12 h light / 12 h dark cycle with *ad libitum* access to standard chow diet (irradiated PicoLab Rodent Diet 20 – 5053 or SDS CRMp). We have maintained the line on a C57BL/6J background for more than twenty generations. We use the terminology ‘knock-out’ (KO) for homozygous mutant mice. We have bred the animals and performed experimental work under licence (PP0116966) issued by the Home Office (UK) in accordance with the Animal (Scientific Procedures) Act 1986 and approved by the Animal Welfare and Ethical Review Body of the University of Liverpool. We are reporting our animal data in line with the ARRIVE guidelines (Percie du Sert et al., 2020).

### Western blot and antibodies

We lysed tissues in RIPA lysis buffer (Merck Life Sciences, Gillingham, UK) supplemented with protease inhibitor cocktail (Merck Life Sciences). We determined protein concentration with Bradford reagent (Merck Life Sciences). We treated lysate samples (20 µg) with NuPAGE LDS sample buffer and reducing agent (Thermo Fisher Scientific, Loughborough, UK), loaded them onto NuPAGE Bis-Tris polyacrylamide 4-12 % gradient gels (Thermo Fisher Scientific) and run them in NuPAGE MOPS running buffer supplemented with NuPAGE antioxidant. We transferred proteins onto Immobilion IF PDVF membrane (Merck Life Sciences) with NuPAGE transfer buffer (Thermo Fisher Scientific) supplemented with 10 % methanol and antioxidant. We incubated membranes with diluted primary antibodies in Odyssey blocking buffer (LI-COR, Cambridge, UK), washed in PBS-Tween (0.1 %) and incubated them with IRDye 680 and 800 secondary antibodies (LI-COR) in PBS-Tween-SDS (0.1 % and 0.01 %) followed by scanning blots in an Odyssey Imaging System (LI-COR). A rabbit polyclonal antibody for Trappc9 was custom produced against the peptide ‘RVESRPTNPSEGS’ and affinity-purified by Dundee Cell Products (Dundee, UK). The β-Actin antibody was from Abcam (Cambridge, UK).

### RNA extraction and RT-PCR

We extracted total RNA from tissues or cells using RNeasy kits (Qiagen, Manchester, UK) or TRIzol reagent (Thermo Fisher Scientific). We generated cDNA with random hexamer primers and ProtoScript II Reverse Transcriptase (New England Biolabs, Hitchin, UK) or SuperScript III Reverse Transcriptase (Thermo Fisher Scientific). We performed PCR with GoTaq Hot Start Polymerase (Promega, Southampton, UK) or Q5 Hot Start High-Fidelity DNA Polymerase (New England Biolabs).

### Histology, immunohistochemistry and *in situ* hybridisation

We dissected tissues for histological analyses from perfusion-fixed mice followed by additional fixation in 4 % PFA/PBS overnight. We dehydrated brain tissues in 30 % sucrose/PBS and prepared 12 μm cryostat sections. For immunohistochemistry with Sox2 antibody (R&D Systems, Abingdon, UK) we incubated sections in 10 mM Na-citrate buffer at 65°C for 5 min for antigen retrieval, quenched endogenous peroxidase activity in methanol / 0.3 % H_2_O_2_, washed in PBS, blocked in PBS / 10 % serum / 0.25 % Triton-X100, incubated with primary antibody at 4°C over night, followed by Vectastain Elite ABC kit (Vector Laboratories, Newark, USA) HRP signal detection, dehydration and embedding in Eukitt mounting medium (Merck Life Sciences). Sox2-positive cells were counted within the demarcated area of the anterior dentate gyrus of matched wild-type (WT) and KO sections (corresponding to Paxinos and Franklin mouse brain atlas plates 45-49) (Paxinos & Franklin, 2001) (Figure 4B) from three WT (13 section averages) and three KO brains (13 section averages). We stained for *Trappc9* RNA expression using the RNAscope 2.5 HD Reagent Kit - RED or RNAscope 2.5 HD Duplex Reagent kit (Advanced Cell Diagnostics, Abingdon, UK) following the manufacturer’s instructions for temperature and incubation times. Prior to staining, we blocked the activity of endogenous peroxidases using RNAscope Hydrogen Peroxide solution for 10 min at room temperature, followed by antigen retrieval via boiling sections in RNAscope Target Retrieval solution for 5 min, washing with distilled water and 100 % ethanol and air-drying. We treated sections with RNAscope Protease Plus solution before incubating them with RNAscope probes Mm-Trappc9-E1E2, Mm-Sox2-C2 and RNAscope negative control probe (Advanced Cell Diagnostics). We washed sections with RNAscope Wash Buffer and incubated them with Hybridize AMP reagents according to the kit protocol. We counterstained the sections with haematoxylin or directly mounted them with Ecomount (Biocare Medical, Pacheco, USA) after air-drying and dipping in Histo-Clear II (National Diagnostics, Atlanta, USA). For adipose tissue histology, we embedded fixed inguinal white and interscapular brown adipose tissues in paraffin, sectioned them at 7 µm thickness, stained with haematoxylin-eosin (H&E) and embedded them in Eukitt.

### MRI data acquisition and processing

We performed all MRI experiments on a 9.4 Tesla horizontal bore magnet USR20 with a Bruker Paravision Console (Bruker Biospin, Bruker, Coventry, UK). For newborn (P0) and weaning ages (P23-P27) we performed MRI *ex vivo* on whole heads that were stored in 4% PFA/PBS. In order to reduce the T1 relaxation time and enhance image contrast, we immersed the samples in a 10 mM solution of the gadolinium-DTPA contrast agent Multihance (Bracco, Milan, Italy) for 24 hrs (newborn) or 72 hrs (weaning age) prior to imaging. We placed samples in custom-made plastic holders that were filled with a fluorinated oil (Fomblin, SolvaySolexis, Brussels, Belgium) to avoid background signal. We performed imaging using a 27 mm loop gap resonator transmit-receive coil (PulseTeq, Chobham, UK). We acquired T1-weighted images using a 3D Fast Low Angle Shot (FLASH) sequence with the parameters: TE: 8 ms, TR: 70.8 ms, Averages: 2, Flip Angle: 60°, Field of View: 12 mm^3^, isotropic resolution: 0.05 mm^3^, Acquisition Time: 2 hrs 33 minutes.

For longitudinal *in vivo* MRI of mice between 4 – 10 months of age we used 1.5 – 2 % isoflurane inhalation anaesthesia. We used a rectal probe to monitor body temperature and a respiratory pillow to measure respiration rate, which we maintained at 50-60 breathes/minute by adjusting the isoflurane level as needed. We placed the mice into a specially designed plastic bed with a face mask fitted to maintain the supply of isoflurane, and ear bars applied to minimise head movement. We placed a heating blanket with circulating warm water over the animal body to maintain body temperature between 31-35°C during the scan. We fitted a 4-channel phased array receiver coil over the head of the mice and the probe, including the 4-channel receive coil, was inserted in a 86 mm transmit coil (Bruker). After running a localizer sequence to generate scout images and adjusting basic frequency, reference power, receiver gain and shims, we used T2-weighted Rapid Imaging with Refocused Echoes (RARE) sequence for anatomical scans covering the whole head with the scanning parameters: TE: 33 ms, TR: 3200 ms, Averages: 8, Echo Spacing: 11 ms, Rare Factor: 8, Field of View: 18 mm^2^, Slices: 30, Slice Thickness: 0.5 mm, Acquisition Matrix: 256 mm^2^, Acquisition Time: 14 min.

DTI was performed using the same field of view and slices as the T2 weighted images above using a spin echo, segmented echo planar imaging (EPI) sequence with the parameters: TE: 22 ms, TR: 3000 ms, Averages: 4, Segments: 6, Directions: 30, Field of View: 18 mm^2^, Slices: 30, Slice Thickness: 0.5 mm, Acquisition Matrix: 96 mm^2^, Acquisition Time: 42 min. In addition, field of view saturation bands were used to cover the ears and areas outside of the brain to reduce inhomogeneity artefacts from these regions. For the DTI scans, respiratory gating was used with the signal from the respiratory monitor (Small Animal Instruments, Stony Brook, USA). Signal acquisition was adjusted such that acquisition only took place during the exhale phase, reducing motion artefacts.

For image processing and analysis, we downloaded files from the MRI Scanner through the ParaVision 6.0.1 software (Bruker) in Digital Imaging and Communication in Medicine (DICOM) format and converted them into the NeuroImaging Information Technology Initiative (NifTI) format with the open source software MRICron. For the determination of total brain volumes, we manually segmented the contrast-enhanced T1-weighted images from *ex vivo* scans, as well as T2-weighted *in vivo* images using the Amira software (Stalling et al., 2005). We constructed the brain segmentation masks using a combination of thresholding tools and manually drawing the masks onto the images using the semi-automated ‘lasso’ and fully manual paintbrush tools. The interpolation tool allowed the segmentation of every other slice and then filling in the alternate slices automatically. We corrected small mistakes made by the program in these slices until the whole brain was covered by the mask.

For the determination of brain sub-region volumes, we used 38 WT and 45 KO littermates at age 4-months. We applied automated segmentation with Atlas-Based Image Data Analysis for Magnetic Resonance Imaging (AIDAmri) (Pallast et al., 2019), which can segment the mouse brain into regions based on the Allen Reference Atlas, on a virtual Linux VMWare Workstation 16. We re-oriented, bias field corrected and skull stripped the NifTI files using a combination of python scripts and FSL commands (Jenkinson et al., 2012). For the extraction of volumes of interest, we registered the images with the Allen Reference Atlas. We overlaid the atlas files over the brain (Figure 3-figure supplement 1A), extracted T2-weighted images in ITK-SNAP (Yushkevich et al., 2006) and extracted regional volumes into .csv files. We then created a searchable database of volumes for each animal brain in Excel using the Database Query function. The process was automated using bash scripts.

We constructed maps from the DTI images after conversion to Nifti. We chose FSL (Jenkinson et al., 2012) to run in the same virtual machine that AIDA was installed on. We wrote a script in MATLAB that searched the Bruker methods file from the original DICOM and extracted the b value/vector files in a format compatible with FSL. Using these files, we followed the DTIFIT pipeline in FSL, which was done using the FSL GUI. FSL models and outputs a number of parametric maps for each image. To improve the reliability of segmentation, we registered the A0 images together by FSLs FLIRT tool. We chose one image as a reference and registered all other images into that space. Since the parametric maps were in the same space as the A0 images, we were able to apply the transforms from this step to all of the individual parameter maps to co-register with each other. We carried out the segmentation of the corpus callosum and the cerebellar white matter in ITK-SNAP (Yushkevich et al., 2006). To get a better picture of the axonal organisation across the corpus callosum, we segmented it into three areas: the genu, towards the anterior, the isthmus near the centre and the splenium at the posterior of the corpus callosum. While these images were aligned well enough in the Z-plane, manual adjustments were occasionally needed where variations in neuroanatomy and slight shifts in the X-Y plane required them. The variation in the shape and size of the cerebellar white matter, despite the registration, made segmenting the full cerebellum for each animal unfeasible. Instead, we placed a 2x2x2 voxel in each hemisphere at the centre of the arbor vitae, adjacent to the parafloculus and 4^th^ ventricle. Examples of the segmentation are shown in Figure 3-figure supplement 1B.

### Behavioural analyses

We performed all tests during the evening (19:00 - 00:00 h) in the dark (active) phase of the animal’s circadian rhythm using the same cohort of WT and KO littermates of both sexes (at age 4 – 6 months) that also underwent *in vivo* MRI. We tested WT and KO littermates in the same experimental session. We undertook the experiments in random consecutive sessions as littermate groups of mice passed through the test age. We cleaned all test equipment with 70 % alcohol and thoroughly dried between trials of each animal. We handled animals daily for a few minutes each for a week before test trials began to accustom them to the experimenter. We performed all animal handling using a tube or by scooping and we never lifted mice by the tail during assays to avoid stress that could influence behaviour (Gouveia & Hurst, 2013; Hurst & West, 2010). We analysed video recordings of animal behaviour and processed the initial data such that they were blinded for the genotype.

The open field apparatus consisted of an open box (92 x 92 cm) with a brightly illuminated (> 1000 lux) centre. We always placed the box in the same position within the room and recorded mouse activity via a camera. Before the trials, we let the mice habituate to the experimental room in their home cages. We always placed the mice into the same corner of the open field box. The mouse was free to explore the open field for 10 minutes in the absence of the experimenter. We analysed the video files with SMART 3.0.1 software (Panlab, Harvard Apparatus, Waterbeach, UK). We used a still of the video to calibrate the software for distance measurements and definition of a 4 x 4 grid of zones. The centre four squares constituted the centre of the field and the remaining squares the periphery. The detection settings of the software fell in a range of Detection Threshold: 6 - 10 and Erosion: 8 - 16 (arbitrary units). The software analysed the video and the experimenter ensured no errors occurred in the automated detection and tracking.

For the accelerating rotarod test we used a Harvard Apparatus Panlab Rota-rod R8 instrument, which automatically recorded the time and speed of rotation for each fall. We carried out the rotarod tests under artificial light (lux range 600-700). We gave mice three training trials per day for 2 days, followed by a third day with two training trials and one test trial (trial 9) (Mann & Chesselet, 2015). We allowed the mice to adjust to the testing room in their home cages for 5 min before placing them on the bar facing against the direction of rotation. We set the rotarod to reach a maximal speed of 44 rpm in a 5 min window. We allowed mice 10 - 30 sec at the start of every trial to get their balance on the bar at its lowest rotation speed before acceleration started, depending on the order, in which mice were placed on the bar. We did not count falls in this time window as part of the trial unless a mouse fell over 3 times, in which case the trial was over and a score of 10 sec was given for time and 4 rpm for acceleration speed. If a mouse had more than 3 of these cases in the first two days (6 trials), we marked them as refusing the task and removed them from the study. After falling from the bar, we rested mice for at least 2 minutes before beginning the next trial. We tested mice from the same home cage only in parallel on different positions of the bar to prevent distraction by unfamiliar scents. We tested a maximum of three mice at a time. The Plug Puzzle is regarded as a test for cognitive problem solving and memory. We made the plug puzzle box according to previously described dimensions (O’Connor et al., 2014) with walls 25 cm high, an open area of 60 x 28 cm, separated from a dark goal box of 28 x 15 cm by a wall with a 4 x 4 cm doorway cut into it. We constructed a U-shaped tunnel of 12 cm length and a wall height of 3.5 cm, which we placed in front of the doorway to the dark box. We blocked the doorway or tunnel escape with different plug materials that offered various degrees of resistance for the mice to overcome without being impossible to remove when trying to escape into the dark box (O’Connor et al., 2014). The first type of blocking material was standard cage bedding, which we used to fill the tunnel. For the second type of blocking material, we used laboratory blue roll tissue plugs, which we formed into a tight ball in a standardised and consistent way. As a third material, we used foam plugs made from closed-cell polyethylene foam designed to protect fragile items in packaging. This foam was durable, moisture resistant and created plugs that did not catch on the doorway and could be removed by continual pushing/pulling on one spot. We gave the mice 3 trials / day for four days with the 5^th^ day featuring as a final test trial (O’Connor et al., 2014). On day 1, we gave the mice a habituation trial (trial 0) with no obstacle blocking the doorway. Then, trials 1 - 3 required the mouse to enter the small tunnel to escape, which was then filled with bedding material for trials 4 - 6. Trial 7 - 9 required the mouse to remove the tissue plug from the doorway, followed by trials 10 – 12, which required the mouse to remove the foam plug from the doorway to escape.

The Novel Object Recognition test took place in the above-mentioned open field box under dim light with camera recording. Based on protocols described previously (Leger et al., 2013), we chose the following objects: a 8 x 3 x 4 cm stack of bricks similar to the well-known ‘Lego’ Brand and cell culture flasks filled half-full with sand, both of an appropriate size for exploration by mice and well balanced to ensure the objects did not fall over during test periods. We prepared these in duplicates to be used in both, the familiarity sessions and the test trials. We allowed the mice to freely explore the empty field for 2 minutes during a short habituation period, since they had already been exposed to the test room and open field box previously. For the familiarisation session on the following day, we chose the type of object (brick stacks or flasks) randomly and placed two identical copies of the object into the field, each 16 cm away from the two opposing walls. We placed the mouse half-way along a perpendicular wall facing the objects. Getting close to the object and sniffing, touching with whiskers and touching with paws counted as active exploration. Climbing on and sitting on the objects did not count as active exploration unless accompanied by sniffing / whisker touches. The probe trial followed a similar structure to the familiarisation trial, but we now replaced one of the familiar objects with a novel object. We randomised the position of the novel object in relation to the mouse (on the left or right) in each trial to avoid any biases. Due to the subjective nature of “active exploration”, which cannot be inferred from proximity to the object alone, we found the captured video files were not suitable for automated analysis as the Open Field test was. Instead, we manually analysed the videos with instances of exploration first watched and noted at full speed, and then the timings of each instance of exploration measured to the nearest frame. During the familiarisation session and probe trial each mouse was given 10 minutes in the test field. We recorded the time spent actively exploring each object. In the probe trial, we analysed for the time taken to reach 20 seconds of overall object exploration (both objects), as well as how much of the 20 seconds exploration time was spent with the novel object.

### Blood plasma analysis for glucose and leptin

We collected blood from ad-libitum fed 9 months-old mice through cardiac puncture and centrifugation at 2000 g for 5 minutes at 4 C° in heparinised tubes (Microvette CB 300; Sarstedt, Leicester, UK) for plasma collection. We measured plasma glucose via a clinical glucose meter (Bayer Contour XT, Bayer Health Care, Leverkusen, Germany). We measured Leptin levels using mouse leptin ELISA kit (Merck Life Sciences) according to manufacturer’s instructions.

### Primary neuron culture

We cultured primary hippocampal neurons as described previously (Beaudoin et al., 2012; Ioannou, Liu, et al., 2019), since *Trappc9* is highly expressed in this brain region. We dissected hippocampi from newborn mouse brain in ice-cold dissection media (HBSS (Merck Life Sciences) supplemented with 0.1 % w/v glucose, 10 mM Hepes pH 7.4, and 1 mM Na-pyruvate) and dissociated them by adding an equal volume of 2 x Papain stock solution (Worthington Biochemical Corporation, Lakewood, USA) at 37°C for 20 minutes. We removed the supernatant carefully and gently rinsed the tissue with plating medium (MEM (Thermo Fisher Scientific) supplemented with 0.45 % glucose, 10 % FBS, 1 mM Na-pyruvate, 2 mM Glutamine, 100 U/ml penicillin, and 0.1 mg/ml streptomycin) before triturating it in fresh plating medium. We filtered the dissociated tissue through a Corning 70 µm cell strainer and collected the cells by centrifugation (200 g, 5 min). We resuspended the cells in neuronal medium (Neurobasal medium (Thermo Fisher Scientific) supplemented with 2 mM glutamine, 100 U/ml penicillin, 0.1 mg/ml streptomycin, 1 x B27 (Thermo Fisher Scientific)) and plated them at a density of 60,000 cells / cm^2^ on KOH-treated, Poly-L-Lysine (Merck Life Sciences) coated coverslips as described (Ioannou, Liu, et al., 2019). We replenished the medium the next day and started selection against replicating non-neuronal cells on day two using neuronal medium containing 2 μM Cytosine β-D- arabinofuranoside (AraC) (Merck Life Sciences). We gradually diluted AraC out by replacing half of the medium with fresh neuronal medium every other day. We used the neurons for experiments from day seven onwards.

### Lipid droplet assay and immunofluorescence imaging

We treated hippocampal neurons with 200 µM Oleic Acid / 0.5 % (w/v) Bovine Serum Albumin (BSA) (HyClone) (premix molar ratio = 2.67/1). We fixed the cells with 4% Paraformaldehyde (PFA) after 6 and 12 hrs of adding OA and incubated with primary and secondary antibodies in PBS, 10 % donkey serum, 0.1 % Saponin followed by staining with LipidTOX (1:1000) (Thermo Fisher Scientific) for 30 minutes and mounting with Vectashield antifade mounting medium (Vector Laboratories). We used a Zeiss LSM800 confocal microscope and acquired images using a Z-stack with 0.5 μm interval. We processed the images with Imaris software (version 9.9) to perform 3D structure analysis. We only analysed lipid droplets located within the cell bodies of neurons to avoid ambiguities in attributing LDs that were located in entangled neurites to specific neurons. For Plin2 analysis, we processed images first to decrease the signal/noise ratio using baseline subtraction. We used the surface generation wizard to generate a surface object for lipid droplets and Plin2 with the parameters provided in the supplementary methods.

### Statistics

We analysed categorical data using Fisher’s Exact or Chi-Square tests. For numerical data we used Student’s paired or independent two-tailed *t.*test, two-way or repeated-measures ANOVA with Šídák’s multiple comparison test or, if data distribution was skewed, Mann-Whitney U-test. Details of statistical tests used for specific datasets are provided in the figure legends, tables or main text. We analysed data with GraphPad Prism software (version 9.3). We considered a *p*-value <0.05 as significant.

### Data availability

All data analysed during this study are included in the manuscript, source data files and figure supplements. Original MRI acquisition files, immunohistochemistry or confocal cell images can be provided upon request.

## Acknowledgements

We would like to thank the University of Liverpool Centre for Cell Imaging, especially Dr Marie Held and Dr Marco Marcello for excellent support in cell imaging and image analysis techniques, Liverpool University Biobank for adipose tissue sectioning and the Biomedical Services Unit for expert animal management and provision of behavioural testing equipment. Furthermore, we would like to thank Dr Mahon Maguire of the Centre for Pre-clinical Imaging for expert assistance with MATLAB scripts for MRI data analysis.

## Figure supplement legends

**Figure 1–figure supplement 1**. **A)** Western blot for Trappc9 on WT and homozygous KO brain tissue using a custom-made antibody. **B)** A cryptic splice site within the Engrailed2 (En2) part of the tm1a gene-trap cassette disrupts β-Galactosidase (LacZ) expression from the *Trappc9* locus. RT-PCR on brain cDNA with primers spanning the indicated exons of *Trappc9* or primers testing for splicing onto the LacZ gene-trap cassette, respectively. While no splicing onto LacZ could be detected in KO samples, cryptic amplicons (arrows) containing downstream exons were found. The schematic overview depicts the arrangement of the tm1a allele. Below the schematic, a part of the sequence of the cryptic exon 5 — 11 KO amplicon is shown in alignment to the expected exon 5 — gene-trap spliced sequence, which should encode the LacZ open reading frame. The alignment indicates that in the observed amplicon splicing initially occurs onto the gene-trap, followed by a cryptic splicing out from the En2 part and onto exon 6 of *Trappc9*, resulting in a mutant transcript that does neither encode β-Galactosidase nor functional Trappc9 protein. SA = splice acceptor site; IRES = internal ribosome entry site.

**Figure 3-figure supplement 1. A)** T2-weighted images (left) overlaid with the Allen Reference Atlas (centre) with a 3D rendering of the whole brain and a dotted line showing the position of the slices (right). **B)** Example A0 image from the DTI image (left) at the splenium of the corpus callosum (top) and cerebellum (bottom). Segmentation maps are shown (right) with the medial and lateral portions of the corpus callosum shown in red and yellow and the arbor vitae voxel in blue.

**Figure 8-figure supplement 1: Adult *Trappc9* KO mice have enlarged lipid droplets in adipose tissues.** Haematoxylin & Eosin stained sections of **A)** white adipose tissue (WAT) and **B)** brown adipose tissue (BAT). Larger adipocytes and lipid droplets were observed in KO tissues compared to WT in both sexes. Scale bar: 100 µm. **C)** RT-PCR from adult mouse adipose tissues confirmed *Trappc9* expression in WAT and BAT. Primers are located in exons 2 and 5.

## Source data file titles

**Figure 1–figure supplement 1–source data 1**. Original Western blot and RT-PCR gel images.

**Figure 2 and Table 1-source data 1**. Raw data for brain volumes measured by MRI.

**Figure 2 and Table 2-source data 2**. Raw data for brain tissue weights.

**Figure 3 and Table 3-source data 1**. Raw data for brain sub-region volumes and DTI.

**Figure 4-source data 1**. Raw data for Sox2-positive NSPCs in dentate gyrus.

**Figure 5 and Table 4-source data 1**. Raw data for all behavioural tests.

**Figure 6-source data 1.** Raw data for lipid droplet volumes per cell and individual LD volumes in hippocampal neurons.

**Figure 7-source data 1.** Raw data for Plin2 positive and negative LDs and Plin2 LD overlap ratios.

**Figure 8-source data 1.** Raw data for body weights, blood glucose and leptin levels.

**Figure 8-figure supplement 1-source data 1**. Original RT-PCR gel images.

## References

1. Al-Deri, N., Okur, V., Ahimaz, P., Milev, M., Valivullah, Z., Hagen, J., Sheng, Y., Chung, W., Sacher, M., & Ganapathi, M. (2021). A novel homozygous variant in TRAPPC2L results in a neurodevelopmental disorder and disrupts TRAPP complex function. J Med Genet, 58(9), 592–601. 10.1136/jmedgenet-2020-107016

2. Almousa, H., Lewis, S. A., Bakhtiari, S., Nordlie, S. H., Pagnozzi, A., Magee, H., Efthymiou, S., Heim, J. A., Cornejo, P., Zaki, M. S., Anwar, N., Maqbool, S., Rahman, F., Neilson, D. E., Vemuri, A., Jin, S. C., Yang, X. R., Heidari, A., van Gassen, K., … Kruer, M. C. (2023). TRAPPC6B biallelic variants cause a neurodevelopmental disorder with TRAPP II and trafficking disruptions. Brain. 10.1093/brain/awad301

3. Amin, M., Vignal, C., Eltaraifee, E., Mohammed, I. N., Hamed, A. A. A., Elseed, M. A., Babai, A., Elbadi, I., Mustafa, D., Abubaker, R., Mustafa, M., Drunat, S., Elsayed, L. E. O., Ahmed, A. E., Boespflug-Tanguy, O., & Dorboz, I. (2022). A novel homozygous mutation in TRAPPC9 gene causing autosomal recessive non-syndromic intellectual disability. BMC Med Genomics, 15(1), 236. 10.1186/s12920-022-01354-1

4. Aslanger, A. D., Goncu, B., Duzenli, O. F., Yucesan, E., Sengenc, E., & Yesil, G. (2022). Biallelic loss of TRAPPC9 function links vesicle trafficking pathway to autosomal recessive intellectual disability. Journal of Human Genetics, 67(5), 279–284. 10.1038/s10038-021-01007-8

5. Beaudoin, G. M., 3rd, Lee, S. H., Singh, D., Yuan, Y., Ng, Y. G., Reichardt, L. F., & Arikkath, J. (2012). Culturing pyramidal neurons from the early postnatal mouse hippocampus and cortex. Nat Protoc, 7(9), 1741–1754. 10.1038/nprot.2012.099

6. Bekbulat, F., Schmitt, D., Feldmann, A., Huesmann, H., Eimer, S., Juretschke, T., Beli, P., Behl, C., & Kern, A. (2020). RAB18 Loss Interferes With Lipid Droplet Catabolism and Provokes Autophagy Network Adaptations. J Mol Biol, 432(4), 1216–1234. 10.1016/j.jmb.2019.12.031

7. Bem, D., Yoshimura, S., Nunes-Bastos, R., Bond, F. C., Kurian, M. A., Rahman, F., Handley, M. T., Hadzhiev, Y., Masood, I., Straatman-Iwanowska, A. A., Cullinane, A. R., McNeill, A., Pasha, S. S., Kirby, G. A., Foster, K., Ahmed, Z., Morton, J. E., Williams, D., Graham, J. M., … Aligianis, I. A. (2011). Loss-of-function mutations in RAB18 cause Warburg micro syndrome. Am J Hum Genet, 88(4), 499–507. 10.1016/j.ajhg.2011.03.012

8. Ben Ayed, I., Bouchaala, W., Bouzid, A., Feki, W., Souissi, A., Ben Nsir, S., Ben Said, M., Sammouda, T., Majdoub, F., Kharrat, I., Kamoun, F., Elloumi, I., Kamoun, H., Tlili, A., Masmoudi, S., & Triki, C. (2021). Further insights into the spectrum phenotype of TRAPPC9 and CDK5RAP2 genes, segregating independently in a large Tunisian family with intellectual disability and microcephaly. Eur J Med Genet, 64(12), 104373. 10.1016/j.ejmg.2021.104373

9. Bolat, H., Unsel-Bolat, G., Derin, H., Sen, A., & Ceylaner, S. (2022). Distinct Autism Spectrum Disorder Phenotype and Hand-Flapping Stereotypes: Two Siblings with Novel Homozygous Mutation in TRAPPC9 Gene and Literature Review. Mol Syndromol, 13(4), 263–269. 10.1159/000522041

10. Bruning, J. C., & Fenselau, H. (2023). Integrative neurocircuits that control metabolism and food intake. Science, 381(6665), eabl7398. 10.1126/science.abl7398

11. Cai, H., Zhang, Y., Pypaert, M., Walker, L., & Ferro-Novick, S. (2005). Mutants in trs120 disrupt traffic from the early endosome to the late Golgi. J Cell Biol, 171(5), 823–833. 10.1083/jcb.200505145

12. Carpanini, S. M., McKie, L., Thomson, D., Wright, A. K., Gordon, S. L., Roche, S. L., Handley, M. T., Morrison, H., Brownstein, D., Wishart, T. M., Cousin, M. A., Gillingwater, T. H., Aligianis, I. A., & Jackson, I. J. (2014). A novel mouse model of Warburg Micro syndrome reveals roles for RAB18 in eye development and organisation of the neuronal cytoskeleton. Dis Model Mech, 7(6), 711–722. 10.1242/dmm.015222

13. Cheng, C. Y., Wu, J. C., Tsai, J. W., Nian, F. S., Wu, P. C., Kao, L. S., Fann, M. J., Tsai, S. J., Liou, Y. J., Tai, C. Y., & Hong, C. J. (2015). ENU mutagenesis identifies mice modeling Warburg Micro Syndrome with sensory axon degeneration caused by a deletion in Rab18. Exp Neurol, 267, 143–151. 10.1016/j.expneurol.2015.03.003

14. Chung, J., Park, J., Lai, Z. W., Lambert, T. J., Richards, R. C., Zhang, J., Walther, T. C., & Farese, R. V., Jr. (2023). The Troyer syndrome protein spartin mediates selective autophagy of lipid droplets. Nat Cell Biol, 25(8), 1101–1110. 10.1038/s41556-023-01178-w

15. Claxton, M., Pulix, M., Seah, M. K. Y., Bernardo, R., Zhou, P., Aljuraysi, S., Liloglou, T., Arnaud, P., Kelsey, G., Messerschmidt, D. M., & Plagge, A. (2022). Variable allelic expression of imprinted genes at the Peg13, Trappc9, Ago2 cluster in single neural cells. Front Cell Dev Biol, 10, 1022422. 10.3389/fcell.2022.1022422

16. Court, F., Camprubi, C., Garcia, C. V., Guillaumet-Adkins, A., Sparago, A., Seruggia, D., Sandoval, J., Esteller, M., Martin-Trujillo, A., Riccio, A., Montoliu, L., & Monk, D. (2014). The PEG13-DMR and brain- specific enhancers dictate imprinted expression within the 8q24 intellectual disability risk locus. Epigenetics Chromatin, 7(1), 5. 10.1186/1756-8935-7-5

17. Deng, Y., Zhou, C., Mirza, A. H., Bamigbade, A. T., Zhang, S., Xu, S., & Liu, P. (2021). Rab18 binds PLIN2 and ACSL3 to mediate lipid droplet dynamics. Biochim Biophys Acta Mol Cell Biol Lipids, 1866(7), 158923. 10.1016/j.bbalip.2021.158923

18. Denoth-Lippuner, A., & Jessberger, S. (2021). Formation and integration of new neurons in the adult hippocampus. Nat Rev Neurosci, 22(4), 223–236. 10.1038/s41583-021-00433-z

19. Ferguson-Smith, A. C. (2011). Genomic imprinting: the emergence of an epigenetic paradigm. Nat Rev Genet, 12(8), 565–575. 10.1038/nrg3032

20. Galindo, A., & Munro, S. (2023). The TRAPP complexes: oligomeric exchange factors that activate the small GTPases Rab1 and Rab11. FEBS Lett, 597(6), 734–749. 10.1002/1873-3468.14553

21. Goncalves, J. T., Schafer, S. T., & Gage, F. H. (2016). Adult Neurogenesis in the Hippocampus: From Stem Cells to Behavior. Cell, 167(4), 897–914. 10.1016/j.cell.2016.10.021

22. Gouveia, K., & Hurst, J. L. (2013). Reducing mouse anxiety during handling: effect of experience with handling tunnels. PLoS One, 8(6), e66401. 10.1371/journal.pone.0066401

23. Hnoonual, A., Graidist, P., Kritsaneepaiboon, S., & Limprasert, P. (2019). Novel Compound Heterozygous Mutations in the TRAPPC9 Gene in Two Siblings With Autism and Intellectual Disability. Front Genet, 10, 61. 10.3389/fgene.2019.00061

24. Hu, M., Bodnar, B., Zhang, Y., Xie, F., Li, F., Li, S., Zhao, J., Zhao, R., Gedupoori, N., Mo, Y., Lin, L., Li, X., Meng, W., Yang, X., Wang, H., Barbe, M. F., Srinivasan, S., Bethea, J. R., Mo, X., … Hu, W. (2023). Defective neurite elongation and branching in Nibp/Trappc9 deficient zebrafish and mice. Int J Biol Sci, 19(10), 3226–3248. 10.7150/ijbs.78489

25. Hurst, J. L., & West, R. S. (2010). Taming anxiety in laboratory mice. Nat Methods, 7(10), 825–826. 10.1038/nmeth.1500

26. Inloes, J. M., Hsu, K. L., Dix, M. M., Viader, A., Masuda, K., Takei, T., Wood, M. R., & Cravatt, B. F. (2014). The hereditary spastic paraplegia-related enzyme DDHD2 is a principal brain triglyceride lipase. Proc Natl Acad Sci U S A, 111(41), 14924–14929. 10.1073/pnas.1413706111

27. Inloes, J. M., Kiosses, W. B., Wang, H., Walther, T. C., Farese, R. V., Jr., & Cravatt, B. F. (2018). Functional Contribution of the Spastic Paraplegia-Related Triglyceride Hydrolase DDHD2 to the Formation and Content of Lipid Droplets. Biochemistry, 57(5), 827–838. 10.1021/acs.biochem.7b01028

28. Ioannou, M. S., Jackson, J., Sheu, S. H., Chang, C. L., Weigel, A. V., Liu, H., Pasolli, H. A., Xu, C. S., Pang, S., Matthies, D., Hess, H. F., Lippincott-Schwartz, J., & Liu, Z. (2019). Neuron-Astrocyte Metabolic Coupling Protects against Activity-Induced Fatty Acid Toxicity. Cell, 177(6), 1522–1535 e1514. 10.1016/j.cell.2019.04.001

29. Ioannou, M. S., Liu, Z., & Lippincott-Schwartz, J. (2019). A Neuron-Glia Co-culture System for Studying Intercellular Lipid Transport. Curr Protoc Cell Biol, 84(1), e95. 10.1002/cpcb.95

30. Jenkins, M. L., Harris, N. J., Dalwadi, U., Fleming, K. D., Ziemianowicz, D. S., Rafiei, A., Martin, E. M., Schriemer, D. C., Yip, C. K., & Burke, J. E. (2020). The substrate specificity of the human TRAPPII complex’s Rab-guanine nucleotide exchange factor activity. Commun Biol, 3(1), 735. 10.1038/s42003-020-01459-2

31. Jenkinson, M., Beckmann, C. F., Behrens, T. E., Woolrich, M. W., & Smith, S. M. (2012). Fsl. Neuroimage, 62(2), 782–790. 10.1016/j.neuroimage.2011.09.015

32. Kandel, P., Semerci, F., Mishra, R., Choi, W., Bajic, A., Baluya, D., Ma, L., Chen, K., Cao, A. C., Phongmekhin, T., Matinyan, N., Jimenez-Panizo, A., Chamakuri, S., Raji, I. O., Chang, L., Fuentes-Prior, P., MacKenzie, K. R., Benn, C. L., Estebanez-Perpina, E., … Maletic-Savatic, M. (2022). Oleic acid is an endogenous ligand of TLX/NR2E1 that triggers hippocampal neurogenesis. Proc Natl Acad Sci U S A, 119(13), e2023784119. 10.1073/pnas.2023784119

33. Ke, Y., Weng, M., Chhetri, G., Usman, M., Li, Y., Yu, Q., Ding, Y., Wang, Z., Wang, X., Sultana, P., DiFiglia, M., & Li, X. (2020). Trappc9 deficiency in mice impairs learning and memory by causing imbalance of dopamine D1 and D2 neurons. Sci Adv, 6(47). 10.1126/sciadv.abb7781

34. Khan, M. A., Khan, S., Windpassinger, C., Badar, M., Nawaz, Z., & Mohammad, R. M. (2016). The Molecular Genetics of Autosomal Recessive Nonsyndromic Intellectual Disability: a Mutational Continuum and Future Recommendations. Ann Hum Genet, 80(6), 342–368. 10.1111/ahg.12176

35. Kiss, R. S., Chicoine, J., Khalil, Y., Sladek, R., Chen, H., Pisaturo, A., Martin, C., Dale, J. D., Brudenell, T. A., Kamath, A., Kyei-Boahen, J., Hafiane, A., Daliah, G., Alecki, C., Hopes, T. S., Heier, M., Aligianis, I. A., Lebrun, J. J., Aspden, J., … Handley, M. T. (2023). Comparative proximity biotinylation implicates the small GTPase RAB18 in sterol mobilization and biosynthesis. J Biol Chem, 299(11), 105295. 10.1016/j.jbc.2023.105295

36. Koifman, A., Feigenbaum, A., Bi, W., Shaffer, L. G., Rosenfeld, J., Blaser, S., & Chitayat, D. (2010). A homozygous deletion of 8q24.3 including the NIBP gene associated with severe developmental delay, dysgenesis of the corpus callosum, and dysmorphic facial features. Am J Med Genet A, 152A(5), 1268–1272. 10.1002/ajmg.a.33319

37. Kramer, J., Beer, M., Bode, H., & Winter, B. (2020). Two Novel Compound Heterozygous Mutations in the TRAPPC9 Gene Reveal a Connection of Non-syndromic Intellectual Disability and Autism Spectrum Disorder. Front Genet, 11, 972. 10.3389/fgene.2020.00972

38. Lamers, I. J. C., Reijnders, M. R. F., Venselaar, H., Kraus, A., Study, D. D. D., Jansen, S., de Vries, B. B. A., Houge, G., Gradek, G. A., Seo, J., Choi, M., Chae, J. H., van der Burgt, I., Pfundt, R., Letteboer, S. J. F., van Beersum, S. E. C., Dusseljee, S., Brunner, H. G., Doherty, D., … Roepman, R. (2017). Recurrent De Novo Mutations Disturbing the GTP/GDP Binding Pocket of RAB11B Cause Intellectual Disability and a Distinctive Brain Phenotype. Am J Hum Genet, 101(5), 824–832. 10.1016/j.ajhg.2017.09.015

39. Leger, M., Quiedeville, A., Bouet, V., Haelewyn, B., Boulouard, M., Schumann-Bard, P., & Freret, T. (2013). Object recognition test in mice. Nat Protoc, 8(12), 2531–2537. 10.1038/nprot.2013.155

40. Li, C., Luo, X., Zhao, S., Siu, G. K., Liang, Y., Chan, H. C., Satoh, A., & Yu, S. S. (2017). COPI-TRAPPII activates Rab18 and regulates its lipid droplet association. EMBO J, 36(4), 441–457. 10.15252/embj.201694866

41. Liang, Z. S., Cimino, I., Yalcin, B., Raghupathy, N., Vancollie, V. E., Ibarra-Soria, X., Firth, H. V., Rimmington, D., Farooqi, I. S., Lelliott, C. J., Munger, S. C., O’Rahilly, S., Ferguson-Smith, A. C., Coll, A. P., & Logan, D. W. (2020). Trappc9 deficiency causes parent-of-origin dependent microcephaly and obesity. PLoS Genet, 16(9), e1008916. 10.1371/journal.pgen.1008916

42. Mann, A., & Chesselet, M. F. (2015). Techniques for Motor Assessment in Rodents. Movement Disorders: Genetics and Models, 2nd Edition, 139–157. 10.1016/B978-0-12-405195-9.00008-1

43. Mir, A., Kaufman, L., Noor, A., Motazacker, M. M., Jamil, T., Azam, M., Kahrizi, K., Rafiq, M. A., Weksberg, R., Nasr, T., Naeem, F., Tzschach, A., Kuss, A. W., Ishak, G. E., Doherty, D., Ropers, H. H., Barkovich, A. J., Najmabadi, H., Ayub, M., & Vincent, J. B. (2009). Identification of mutations in TRAPPC9, which encodes the NIK- and IKK-beta-binding protein, in nonsyndromic autosomal-recessive mental retardation. Am J Hum Genet, 85(6), 909–915. 10.1016/j.ajhg.2009.11.009

44. Mochida, G. H., Mahajnah, M., Hill, A. D., Basel-Vanagaite, L., Gleason, D., Hill, R. S., Bodell, A., Crosier, M., Straussberg, R., & Walsh, C. A. (2009). A truncating mutation of TRAPPC9 is associated with autosomal-recessive intellectual disability and postnatal microcephaly. Am J Hum Genet, 85(6), 897–902. 10.1016/j.ajhg.2009.10.027

45. O’Connor, A. M., Burton, T. J., Leamey, C. A., & Sawatari, A. (2014). The use of the puzzle box as a means of assessing the efficacy of environmental enrichment. J Vis Exp(94). 10.3791/52225

46. Olzmann, J. A., & Carvalho, P. (2019). Dynamics and functions of lipid droplets. Nat Rev Mol Cell Biol, 20(3), 137–155. 10.1038/s41580-018-0085-z

47. Pallast, N., Diedenhofen, M., Blaschke, S., Wieters, F., Wiedermann, D., Hoehn, M., Fink, G. R., & Aswendt, M. (2019). Processing Pipeline for Atlas-Based Imaging Data Analysis of Structural and Functional Mouse Brain MRI (AIDAmri). Front Neuroinform, 13, 42. 10.3389/fninf.2019.00042

48. Paxinos, G., & Franklin, K. B. J. (2001). The mouse brain in stereotaxic coordinates (second edition ed.). Academic Press.

49. Penon-Portmann, M., Hodoglugil, U., Arun, P. W., Yip, T., Slavotinek, A., & Tenney, J. L. (2023). TRAPPC9-related neurodevelopmental disorder: Report of a homozygous deletion in TRAPPC9 due to paternal uniparental isodisomy. Am J Med Genet A, 191(4), 1077–1082. 10.1002/ajmg.a.63100

50. Percie du Sert, N., Hurst, V., Ahluwalia, A., Alam, S., Avey, M. T., Baker, M., Browne, W. J., Clark, A., Cuthill, I. C., Dirnagl, U., Emerson, M., Garner, P., Holgate, S. T., Howells, D. W., Karp, N. A., Lazic, S. E., Lidster, K., MacCallum, C. J., Macleod, M., … Wurbel, H. (2020). The ARRIVE guidelines 2.0: Updated guidelines for reporting animal research. PLoS Biol, 18(7), e3000410. 10.1371/journal.pbio.3000410

51. Perez, J. D., Rubinstein, N. D., Fernandez, D. E., Santoro, S. W., Needleman, L. A., Ho-Shing, O., Choi, J. J., Zirlinger, M., Chen, S. K., Liu, J. S., & Dulac, C. (2015). Quantitative and functional interrogation of parent-of-origin allelic expression biases in the brain. Elife, 4, e07860. 10.7554/eLife.07860

52. Philippe, O., Rio, M., Carioux, A., Plaza, J. M., Guigue, P., Molinari, F., Boddaert, N., Bole-Feysot, C., Nitschke, P., Smahi, A., Munnich, A., & Colleaux, L. (2009). Combination of linkage mapping and microarray-expression analysis identifies NF-kappaB signaling defect as a cause of autosomal-recessive mental retardation [Research Support, Non-U.S. Gov’t]. Am J Hum Genet, 85(6), 903–908. 10.1016/j.ajhg.2009.11.007

53. Radenkovic, S., Martinelli, D., Zhang, Y., Preston, G. J., Maiorana, A., Terracciano, A., Dentici, M. L., Pisaneschi, E., Novelli, A., Ranatunga, W., Ligezka, A. N., Ghesquiere, B., Deyle, D. R., Kozicz, T., Pinto, E. V. F., Witters, P., & Morava, E. (2022). TRAPPC9-CDG: A novel congenital disorder of glycosylation with dysmorphic features and intellectual disability. Genet Med, 24(4), 894–904. 10.1016/j.gim.2021.12.012

54. Ralhan, I., Chang, C. L., Lippincott-Schwartz, J., & Ioannou, M. S. (2021). Lipid droplets in the nervous system. J Cell Biol, 220(7). 10.1083/jcb.202102136

55. Ramosaj, M., Madsen, S., Maillard, V., Scandella, V., Sudria-Lopez, D., Yuizumi, N., Telley, L., & Knobloch, M. (2021). Lipid droplet availability affects neural stem/progenitor cell metabolism and proliferation. Nat Commun, 12(1), 7362. 10.1038/s41467-021-27365-7

56. Rasika, S., Passemard, S., Verloes, A., Gressens, P., & El Ghouzzi, V. (2018). Golgipathies in Neurodevelopment: A New View of Old Defects. Dev Neurosci, 40(5-6), 396–416. 10.1159/000497035

57. Rawlins, L. E., Almousa, H., Khan, S., Collins, S. C., Milev, M. P., Leslie, J., Saint-Dic, D., Khan, V., Hincapie, A. M., Day, J. O., McGavin, L., Rowley, C., Harlalka, G. V., Vancollie, V. E., Ahmad, W., Lelliott, C. J., Gul, A., Yalcin, B., Crosby, A. H., … Baple, E. L. (2022). Biallelic variants in TRAPPC10 cause a microcephalic TRAPPopathy disorder in humans and mice. PLoS Genet, 18(3), e1010114. 10.1371/journal.pgen.1010114

58. Riedel, F., Galindo, A., Muschalik, N., & Munro, S. (2018). The two TRAPP complexes of metazoans have distinct roles and act on different Rab GTPases. J Cell Biol, 217(2), 601–617. 10.1083/jcb.201705068

59. Robinett, C. C., Giansanti, M. G., Gatti, M., & Fuller, M. T. (2009). TRAPPII is required for cleavage furrow ingression and localization of Rab11 in dividing male meiotic cells of Drosophila. J Cell Sci, 122(Pt 24), 4526–4534. 10.1242/jcs.054536

60. Ruf, N., Bahring, S., Galetzka, D., Pliushch, G., Luft, F. C., Nurnberg, P., Haaf, T., Kelsey, G., & Zechner, U. (2007). Sequence-based bioinformatic prediction and QUASEP identify genomic imprinting of the KCNK9 potassium channel gene in mouse and human. Human Molecular Genetics, 16(21), 2591–2599. 10.1093/hmg/ddm216

61. Rybak, K., Steiner, A., Synek, L., Klaeger, S., Kulich, I., Facher, E., Wanner, G., Kuster, B., Zarsky, V., Persson, S., & Assaad, F. F. (2014). Plant cytokinesis is orchestrated by the sequential action of the TRAPPII and exocyst tethering complexes. Dev Cell, 29(5), 607–620. 10.1016/j.devcel.2014.04.029

62. Sacher, M., Shahrzad, N., Kamel, H., & Milev, M. P. (2019). TRAPPopathies: An emerging set of disorders linked to variations in the genes encoding transport protein particle (TRAPP)-associated proteins. Traffic, 20(1), 5–26. 10.1111/tra.12615

63. Skarnes, W. C., Rosen, B., West, A. P., Koutsourakis, M., Bushell, W., Iyer, V., Mujica, A. O., Thomas, M., Harrow, J., Cox, T., Jackson, D., Severin, J., Biggs, P., Fu, J., Nefedov, M., de Jong, P. J., Stewart, A. F., & Bradley, A. (2011). A conditional knockout resource for the genome-wide study of mouse gene function. Nature, 474(7351), 337–342. 10.1038/nature10163

64. Smith, R. J., Dean, W., Konfortova, G., & Kelsey, G. (2003). Identification of novel imprinted genes in a genome-wide screen for maternal methylation. Genome Research, 13(4), 558–569. 10.1101/gr.781503

65. Stalling, D., Westerhoff, M., & Hege, H. C. (2005). Amira: A highly interactive system for visual data analysis. In C. D. Hansen, C. R. J. Johnson, & C. R. Johnson (Eds.), Visualization Handbook (pp. 749–767). Elsevier Inc. 10.1016/B978-012387582-2/50040-X

66. Sztalryd, C., & Brasaemle, D. L. (2017). The perilipin family of lipid droplet proteins: Gatekeepers of intracellular lipolysis. Biochim Biophys Acta Mol Cell Biol Lipids, 1862(10 Pt B), 1221–1232. 10.1016/j.bbalip.2017.07.009

67. Tucci, V., Isles, A. R., Kelsey, G., Ferguson-Smith, A. C., & Erice Imprinting, G. (2019). Genomic Imprinting and Physiological Processes in Mammals. Cell, 176(5), 952–965. 10.1016/j.cell.2019.01.043

68. Usman, M., Li, Y., Ke, Y., Chhetri, G., Islam, M. A., Wang, Z., & Li, X. (2022). Trappc9 Deficiency Impairs the Plasticity of Stem Cells. Int J Mol Sci, 23(9). 10.3390/ijms23094900

69. Van Bergen, N. J., Guo, Y., Al-Deri, N., Lipatova, Z., Stanga, D., Zhao, S., Murtazina, R., Gyurkovska, V., Pehlivan, D., Mitani, T., Gezdirici, A., Antony, J., Collins, F., Willis, M. J. H., Coban Akdemir, Z. H., Liu, P., Punetha, J., Hunter, J. V., Jhangiani, S. N., … Christodoulou, J. (2020). Deficiencies in vesicular transport mediated by TRAPPC4 are associated with severe syndromic intellectual disability. Brain, 143(1), 112–130. 10.1093/brain/awz374

70. Wang, N., Lee, I. J., Rask, G., & Wu, J. Q. (2016). Roles of the TRAPP-II Complex and the Exocyst in Membrane Deposition during Fission Yeast Cytokinesis. PLoS Biol, 14(4), e1002437. 10.1371/journal.pbio.1002437

71. Xu, D., Li, Y., Wu, L., Li, Y., Zhao, D., Yu, J., Huang, T., Ferguson, C., Parton, R. G., Yang, H., & Li, P. (2018). Rab18 promotes lipid droplet (LD) growth by tethering the ER to LDs through SNARE and NRZ interactions. J Cell Biol, 217(3), 975–995. 10.1083/jcb.201704184

72. Yao, Z., van Velthoven, C. T. J., Nguyen, T. N., Goldy, J., Sedeno-Cortes, A. E., Baftizadeh, F., Bertagnolli, D., Casper, T., Chiang, M., Crichton, K., Ding, S. L., Fong, O., Garren, E., Glandon, A., Gouwens, N. W., Gray, J., Graybuck, L. T., Hawrylycz, M. J., Hirschstein, D., … Zeng, H. (2021). A taxonomy of transcriptomic cell types across the isocortex and hippocampal formation. Cell, 184(12), 3222–3241 e3226. 10.1016/j.cell.2021.04.021

73. Yushkevich, P. A., Piven, J., Hazlett, H. C., Smith, R. G., Ho, S., Gee, J. C., & Gerig, G. (2006). User-guided 3D active contour segmentation of anatomical structures: significantly improved efficiency and reliability. Neuroimage, 31(3), 1116–1128. 10.1016/j.neuroimage.2006.01.015

