## Supplement to Fig 1 for "Microcephaly with a disproportionate hippocampal reduction, stem cell loss and neuronal lipid droplet symptoms in *Trappc9* KO mice"

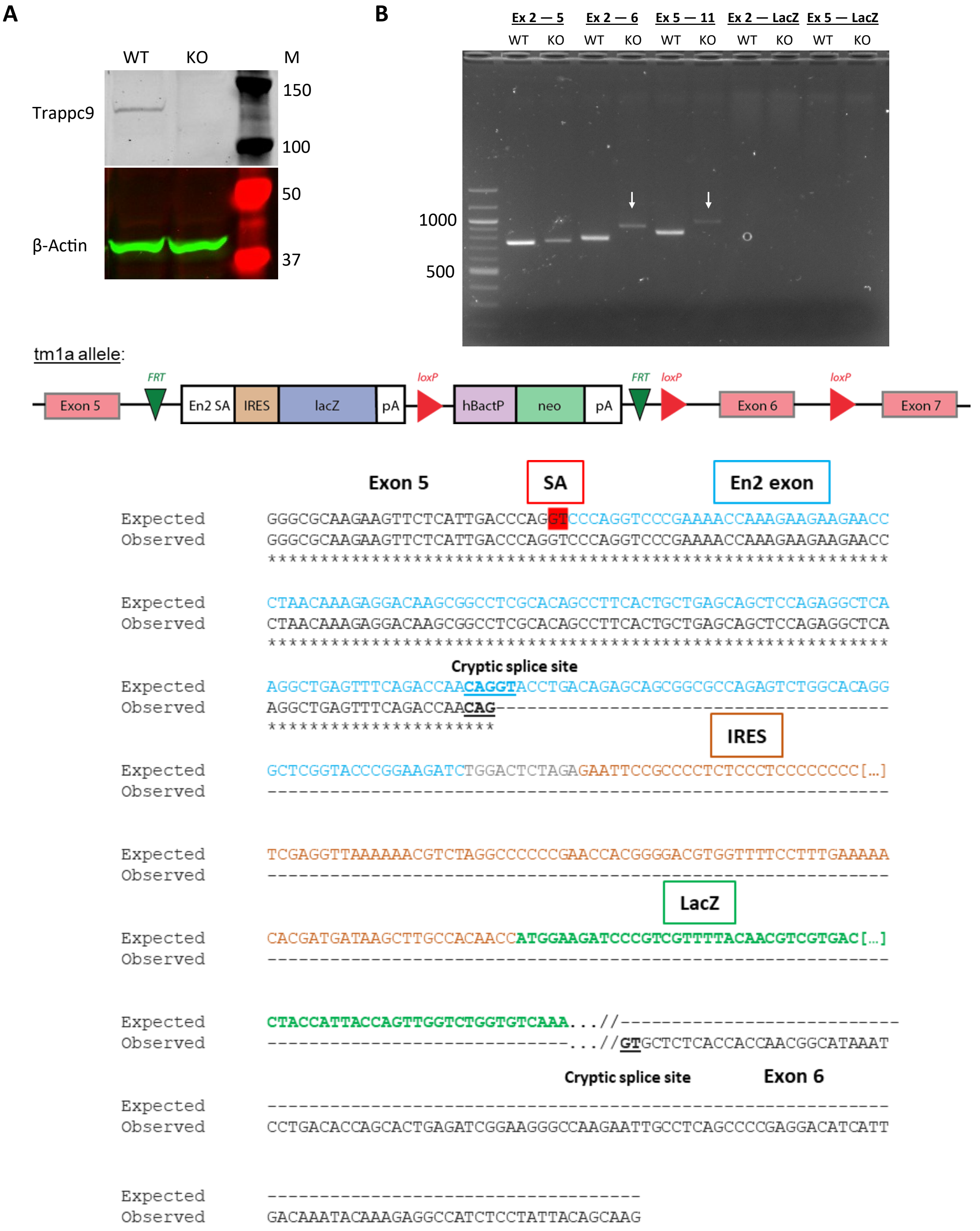


**Figure 1—figure supplement 1. A)** Western blot for Trappc9 on WT and homozygous KO brain tissue using a custom-made antibody. **B)** A cryptic splice site within the Engrailed2 (En2) part of the tm1a gene-trap cassette disrupts β-Galactosidase (LacZ) expression from the *Trappc9* locus. RT-PCR on brain cDNA with primers spanning the indicated exons of *Trappc9* or primers testing for splicing onto the LacZ gene-trap cassette, respectively. While no splicing onto LacZ could be detected in KO samples, cryptic amplicons (arrows) containing downstream exons were found. The schematic overview depicts the arrangement of the tm1a allele. Below the schematic, a part of the sequence of the cryptic exon 5 — 11 KO amplicon is shown in alignment to the expected exon 5 — gene-trap spliced sequence, which should encode the LacZ open reading frame. The alignment indicates that in the observed amplicon splicing initially occurs onto the gene-trap, followed by a cryptic splicing out from the En2 part and onto exon 6 of *Trappc9*, resulting in a mutant transcript that does neither encode β-Galactosidase nor functional Trappc9 protein. SA = splice acceptor site; IRES = internal ribosome entry site.
