## Supplement to Fig 3 for "Microcephaly with a disproportionate hippocampal reduction, stem cell loss and neuronal lipid droplet symptoms in *Trappc9* KO mice"

A

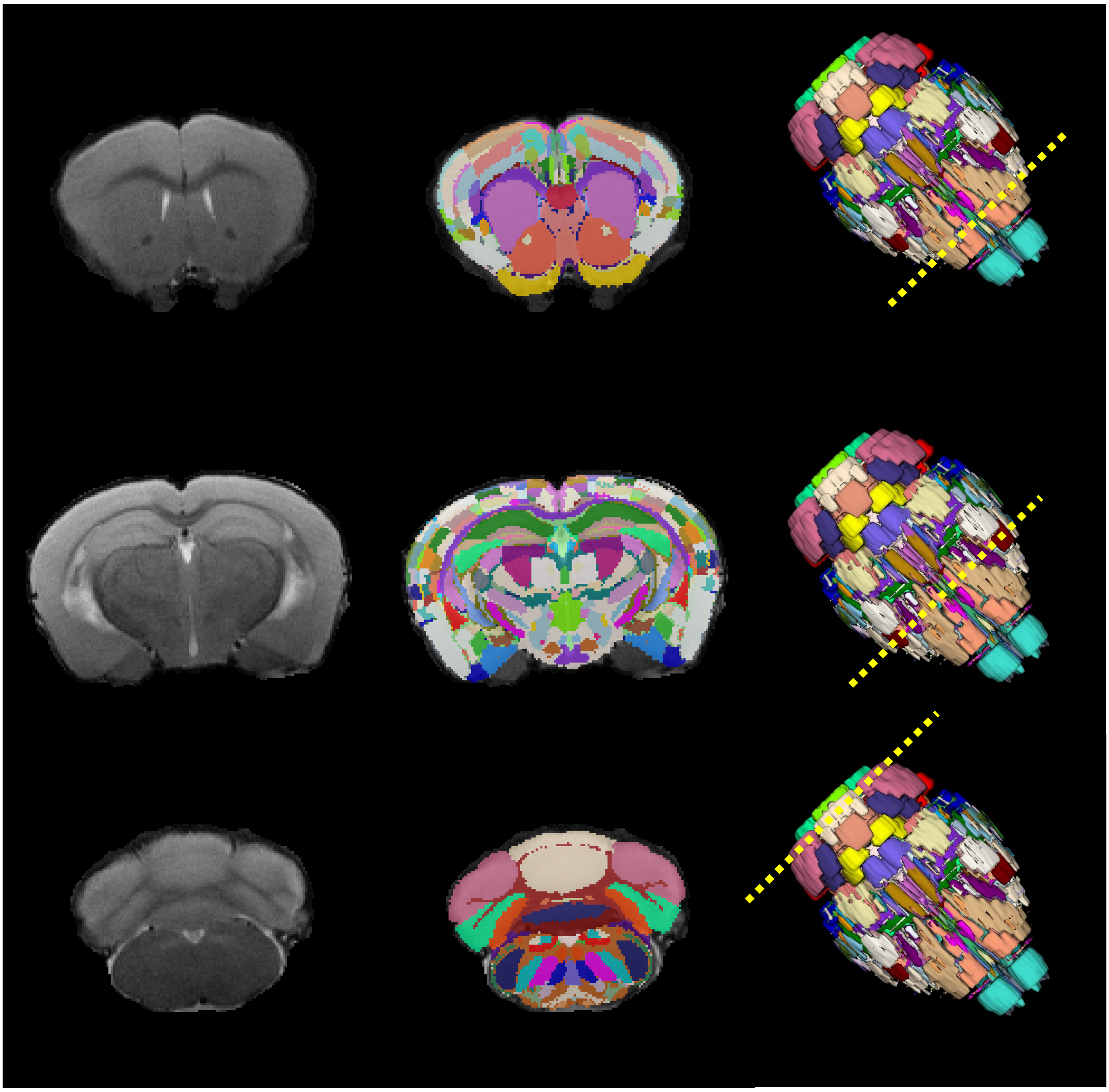

B

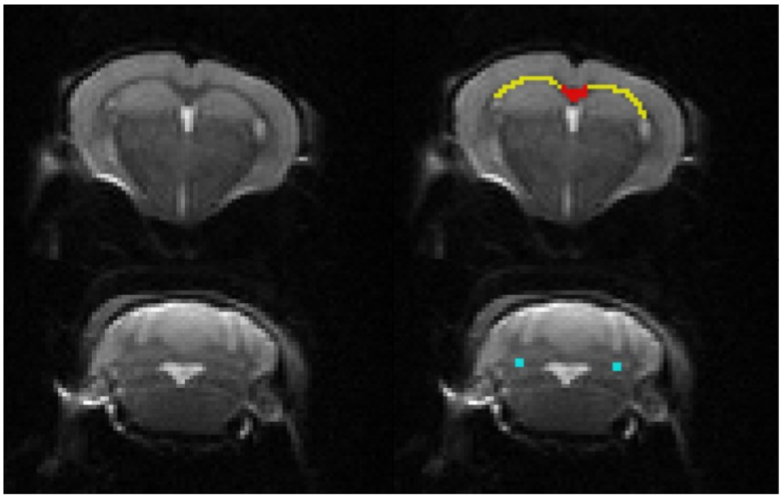

**Figure 3-figure supplement 1. A)** T2-weighted images (left) overlaid with the Allen Reference Atlas (centre) with a 3D rendering of the whole brain and a dotted line showing the position of the slices (right). **B)** Example A0 image from the DTI image (left) at the splenium of the corpus callosum (top) and cerebellum (bottom). Segmentation maps are shown (right) with the medial and lateral portions of the corpus callosum shown in red and yellow and the arbor vitae voxel in blue.
