## Supplement to Fig 8 for "Microcephaly with a disproportionate hippocampal reduction, stem cell loss and neuronal lipid droplet symptoms in *Trappc9* KO mice"

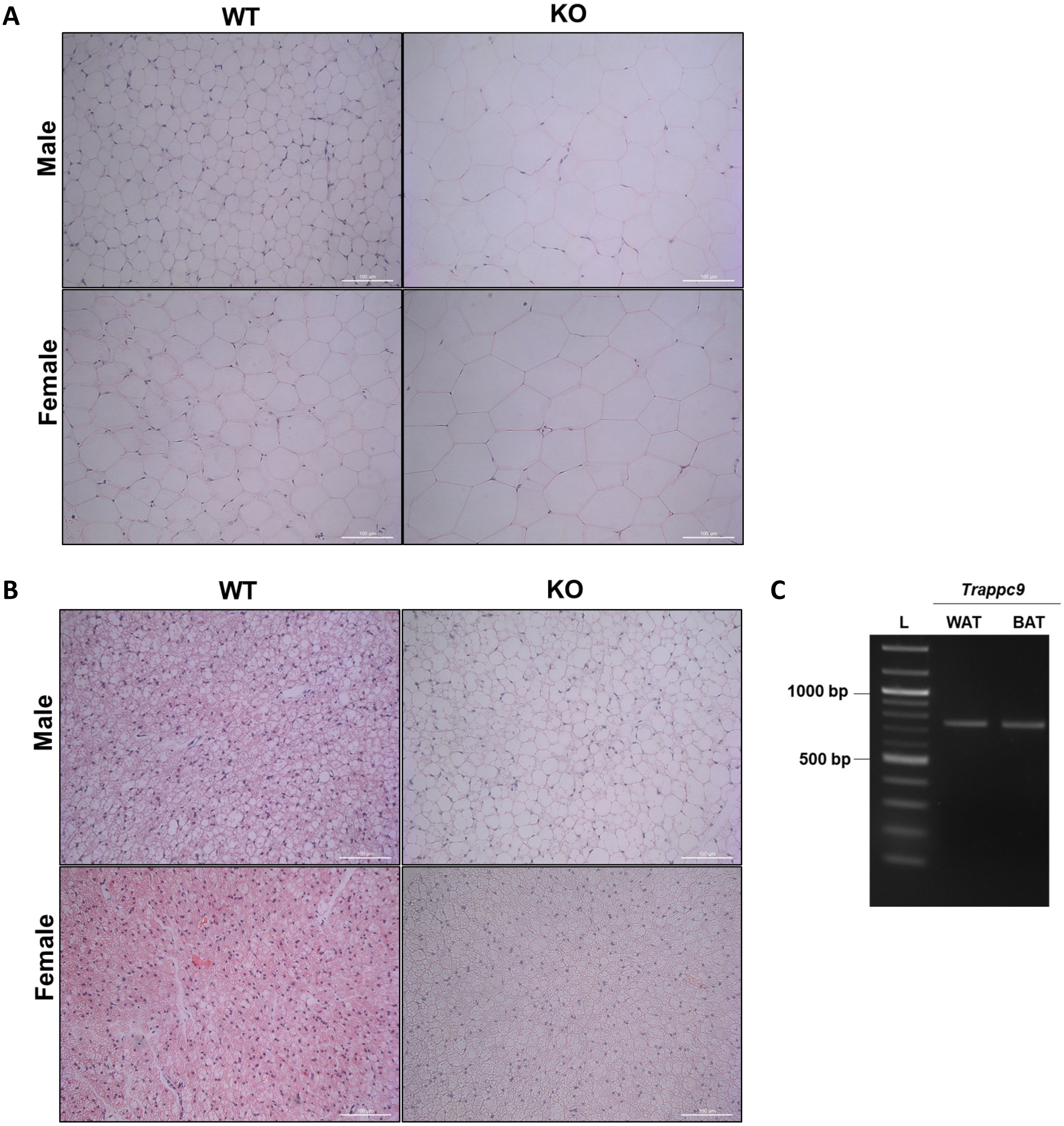


**Figure 8–figure supplement 1: Adult *Trappc9* KO mice have enlarged lipid droplets in adipose tissues.** Haematoxylin & Eosin stained sections of **A)** white adipose tissue (WAT) and **B)** brown adipose tissue (BAT). Larger adipocytes and lipid droplets were observed in KO tissues compared to WT in both sexes. Scale bar: 100 µm. **C)** RT-PCR from adult mouse adipose tissues confirmed *Trappc9* expression in WAT and BAT. Primers are located in exons 2 and 5.
